# Amplicon Sequencing of Single-copy Protein-coding Genes Reveals Accurate Diversity for Sequence-discrete Microbiome Populations

**DOI:** 10.1101/2021.10.22.465537

**Authors:** Chengfeng Yang, Qinzhi Su, Min Tang, Shiqi Luo, Hao Zheng, Xue Zhang, Xin Zhou

## Abstract

An in-depth understanding of microbial function and the division of ecological niches requires accurate delineation and identification of microbes at a fine taxonomic resolution. Microbial phylotypes are typically defined using a 97% small subunit (16S) rRNA threshold. However, increasing evidence has demonstrated the ubiquitous presence of taxonomic units of distinct functions within phylotypes. These so-called sequence-discrete populations (SDPs) have used to be mainly delineated by disjunct sequence similarity at the whole-genome level. However, gene markers that could accurately identify and quantify SDPs are lacking in microbial community studies. Here we developed a pipeline to screen single-copy protein-coding genes that could accurately characterize SDP diversity via amplicon sequencing of microbial communities. Fifteen candidate marker genes were evaluated using three criteria (extent of sequence divergence, phylogenetic accuracy, and conservation of primer regions) and the selected genes were subject to test the efficiency in differentiating SDPs within *Gilliamella*, a core honeybee gut microbial phylotype, as a proof-of-concept. The results showed that the 16S V4 region failed to report accurate SDP diversities due to low taxonomic resolution and changing copy numbers. In contrast, the single-copy genes recommended by our pipeline were able to successfully quantify *Gilliamella* SDPs for both mock samples and honeybee guts, with results highly consistent with those of metagenomics. The pipeline developed in this study is expected to identify single-copy protein coding genes capable of accurately quantifying diverse bacterial communities at the SDP level.

**IMPORTANCE:** Microbial communities can be distinguished by discrete genetic and ecological characteristics. These sequence-discrete populations are foundational for investigating the composition and functional structures of microbial communities at high resolution. In this study, we screened for reliable single-copy protein-coding marker genes to identify sequence-discrete populations through our pipeline. Using marker gene amplicon sequencing, we could accurately and efficiently delineate the population diversity in microbial communities. These results suggest that single copy protein-coding genes can be an accurate, quantitative and economical alternative for characterizing population diversity. Moreover, the feasibility of a gene as marker for any bacterial population identification can be quickly evaluated by the pipeline proposed here.

## INTRODUCTION

Accurate identification of distinct functional units in natural bacterial communities is crucial in understanding their ecological roles, interactions within the network, as well as the fine-scale composition and dynamic changes within the whole community. As a rule of thumb, a bacterial phylotype is often defined by grouping strains that share a sequence identify greater than 97% for a selected fragment of the small subunit (16S) rRNA gene [1]. However, increasing evidence has indicated that a bacterial phylotype may contain multiple finer lineages, each showing distinct biological traits. For example, closely related enterotoxigenic *Escherichia coli* (ETEC) isolates form discrete lineages with consistently definable variations in virulence profiles [2]. Such intra-phylotype lineages could be delineated based on divergence in genomic sequences and phylogenetic inferences. These finer subdivisions of phylotypes are called sequence-discrete populations (SDPs), which typified by genetic and genealogical discontinuity from the rest of the community, and are delineated by overall sequence divergence at the whole-genome level [3–5]. A broad comparison of 90,000 bacterial genomic sequences, with a close examination of pairwise genomic similarities in natural bacterial communities, has proved the pervasive discontinuity in genetic similarity below and above SDPs [3]. Bacteria in the same SDP normally show less than ca. 5% variation in whole-genome sequences. This genetic divergence is much less than those among strains of the same phylotype (ca. 30%) [6]. With respect to habitats, specific SDPs are likely ubiquitous in various environments, such as human and animal guts [5, 7, 8], freshwater [9], ocean [10] and soil [11]. Therefore, SDPs are probably better than phylotypes, as taxonomic units that represent functional entities in bacterial communities, which are likely shaped by ecological pressure and evolutionary selection. As such, SDPs are important units of microbial diversity and should be considered as baseline information for investing crucial questions, such as how do bacterial populations interact and evolve within communities [4].

Despite the essential nature of accurate SDP identification, a rapid and accurate method that can trace SDP boundaries is still lacking, especially with regards to the selection of proper markers for evaluating sequence divergence. It is obvious that genetic divergence among bacterial strains is dependent on which genes are compared. We now understand that the commonly used 16S gene cannot generally provide sufficient resolution to characterize SDP diversity [12, 13]. For example, in cases where the SDPs show a ~5-10% genome-wide divergence, they varied mostly merely < 0.1% in the 16S sequences [14]. Moreover, the copy number of the 16S gene may vary significantly among phylotypes or even among strains of the same phylotype, making quantitative characterization of bacterial community a challenging, if not impossible, task [15, 16]. The 16S was selected for phylotype delineation years ago because it has conserved primer sites that flank relatively variable regions that made it easy to sequence with Sanger technology. Currently, much effort has been put into developing genes or gene segments that can be easily sequenced, and that vary enough to serve as practical proxies for SDP delineation [17–19]. However, a systematic evaluation of the validity and performance of such genes in SDP delineation, which includes the rapidly increasing but heterogeneously sampled database, has not been carried out.

Fortunately, recent developments in microbial genomics show a promising solution to complement the coverage of bacterial genomes. The number of sequenced genomes of various bacterial lineages has been growing rapidly. For example, the Genomes OnLine Database (GOLD) now contains 437,099 bacterial genomes, the majority of which (397,945) are uncultured, representing host-associated, environmental and engineered ecosystems [20]. The ever-growing bacterial genome dataset offers a great opportunity to screen phylogenetically informative genes that show good performance in taxonomic delineation, including those capable of quantitatively charactering bacterial communities at the SDP level [21, 22]. For instance, Wu and colleagues identified 114 PhyEco universal markers for all bacteria [23]. From these universal markers, 15 single-copy protein-coding genes were successfully applied in estimating species abundances using shotgun metagenomic data [24]. On the other hand, growing numbers of genomes and metagenomes produced for particular bacterial communities or taxonomic groups allow for comprehensive characterization of SDP diversity within focal environments and bacterial groups. Taking social bee gut microbiota as an example, diverse strains derived from major honeybee hosts have been isolated and deep-sequenced [25], including well-covered SDPs of nearly all core gut bacterial phylotypes [5, 26, 27]. Thus, the relatively complete genome dataset provides a genome-wide-based gold standard for defining SDPs for the honeybee core bacteria.

In the present study, we developed a pipeline to screen potential marker genes capable of accurate identification and quantification of SDP diversity. We used the core bacterial phylotype *Gilliamella* derived from the eastern honeybee *Apis cerana* as a proof of concept, and delineated *Gilliamella* SDPs based on a set of comprehensive genome sequences. We further screened 15 single-copy protein-coding genes, which are present in all bacteria, to identify candidate marker genes capable of differentiating the defined *Gilliamella* SDPs. Important characteristics such as the level of sequence divergence, phylogenetic robustness, and the presence of conservative primer regions, are considered in marker gene screening. Finally, we applied the candidate markers in amplicon sequencing of both bacterial mock samples and real honeybee guts to verify their efficiency in SDP profiling (Fig. 1). The markers we identified could accurately, consistently and quantitatively capture SDP diversity.

**FIG 1.**
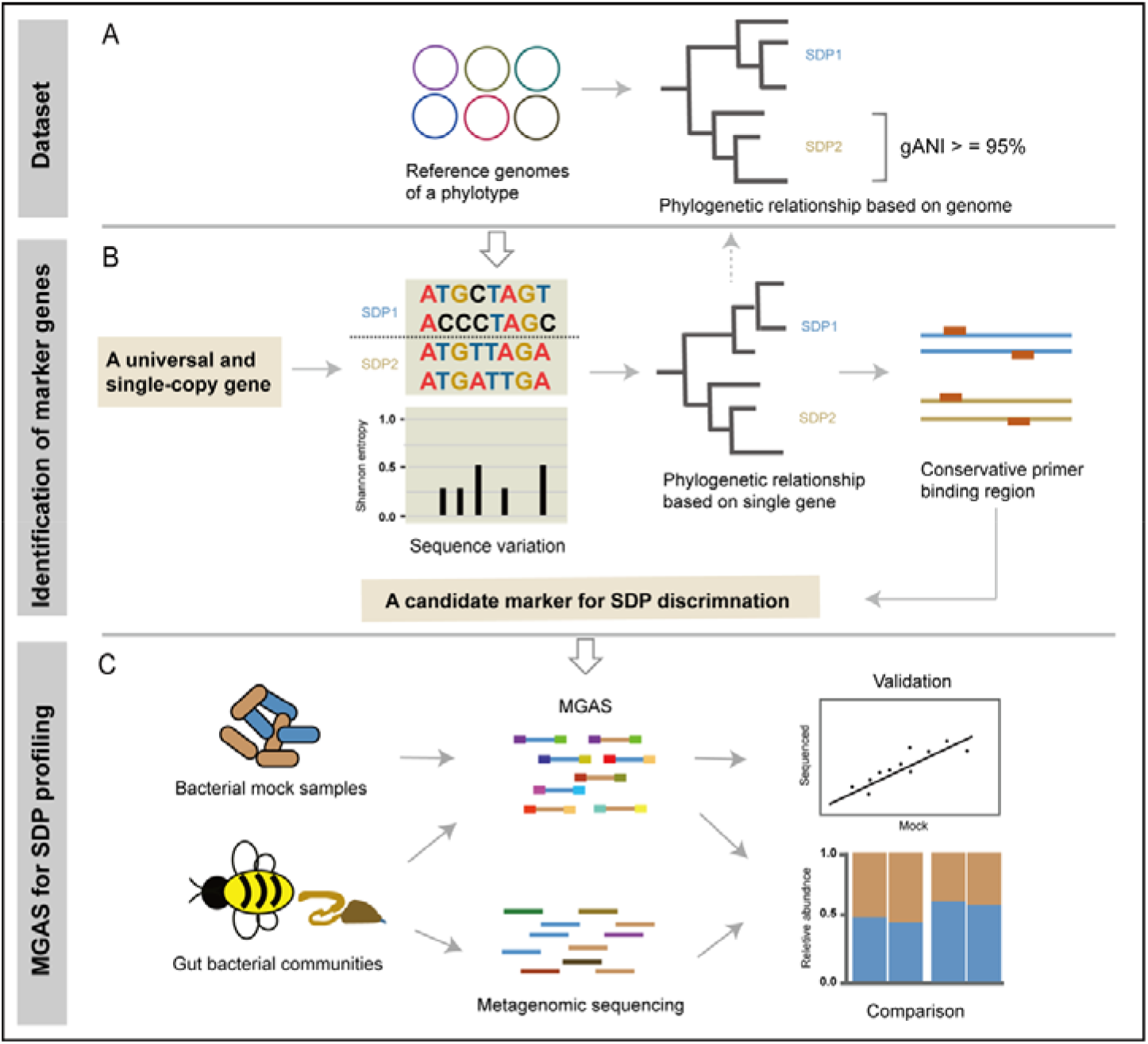
Screening marker genes suitable for SDP discrimination and quantification. (A) SDPs are identified for gut bacterial phylotypes based on phylogenetic relationships and genome-wide pairwise average nucleotide identities (gANI). (B) A candidate marker gene for SDP discrimination is selected from a set of universal and single-copy genes based on sequence variation, phylogenetic relationship and well-conserved regions for primer design. (C) The performance of marker gene amplicon sequencing (MGAS) on SDP identification and quantification is validated and compared as characterized using the mock samples and gut gut communities.

## RESULTS

### A comprehensive genome reference database for honeybee gut bacteria

A comprehensive genome reference database was constructed for honeybee gut bacteria (Table S1). A total of 242 genomes were included, covering 103 isolates from *A. cerana* and 139 from *A. mellifera*. SDPs were identified for the core gut bacterial phylotypes using these reference genomes. SDPs differed between honeybee species, which is consistent with previous studies [27, 28]. Within *A. cerana* phylotypes, 5 SDPs were identified for *Gilliamella* (Gillia, n=65), 2 for *Bifidobacterium* (Bifido, n=9), 1 for *Lactobacillus* Firm5 (Firm5, n=6), 1 for *Apibacter* (Apib, n=16) and 2 for *Snodgrassella* (Snod, n=7). Within *A. mellifera* phylotypes, 6 SDPs were identified for Gillia (n=65), 9 for Bifido (n=19), 2 for *Lactobacillus* Firm4 (Firm4, n=2), 6 for Firm5 (n=18) and 2 for Snod (n=35) (Table S1). These SDPs delineated by genomes were used as references for subsequent taxonomic assignments for the 16S, marker gene, or metagenome-based SDP identifications.

### Single-copy marker genes showed higher sequence variations at the SDP level than the 16S gene

Sufficient sequence variation is crucial for high resolution discrimination of bacterial SDPs. Here we compared the average Shannon entropy (ASE) between the whole-16S and the 15 single-copy marker genes. Our results clearly showed that the marker genes had much higher ASEs at both phylotype and SDP levels compared to those of the 16S (Fig. 2A). The regional difference in the variation levels between 16S and selected marker genes was also compared along the full gene length. A slide-window (20 bp) ASE analysis showed that although several spikes of variable regions were identified along the 16S gene, with the highest variable region corresponded to part of the classic V3 region, its regional ASEs were generally lower compared to marker genes, e.g., *NusA, PTH* and *frr* (Fig. 2B; Fig. S1).

**FIG 2.**
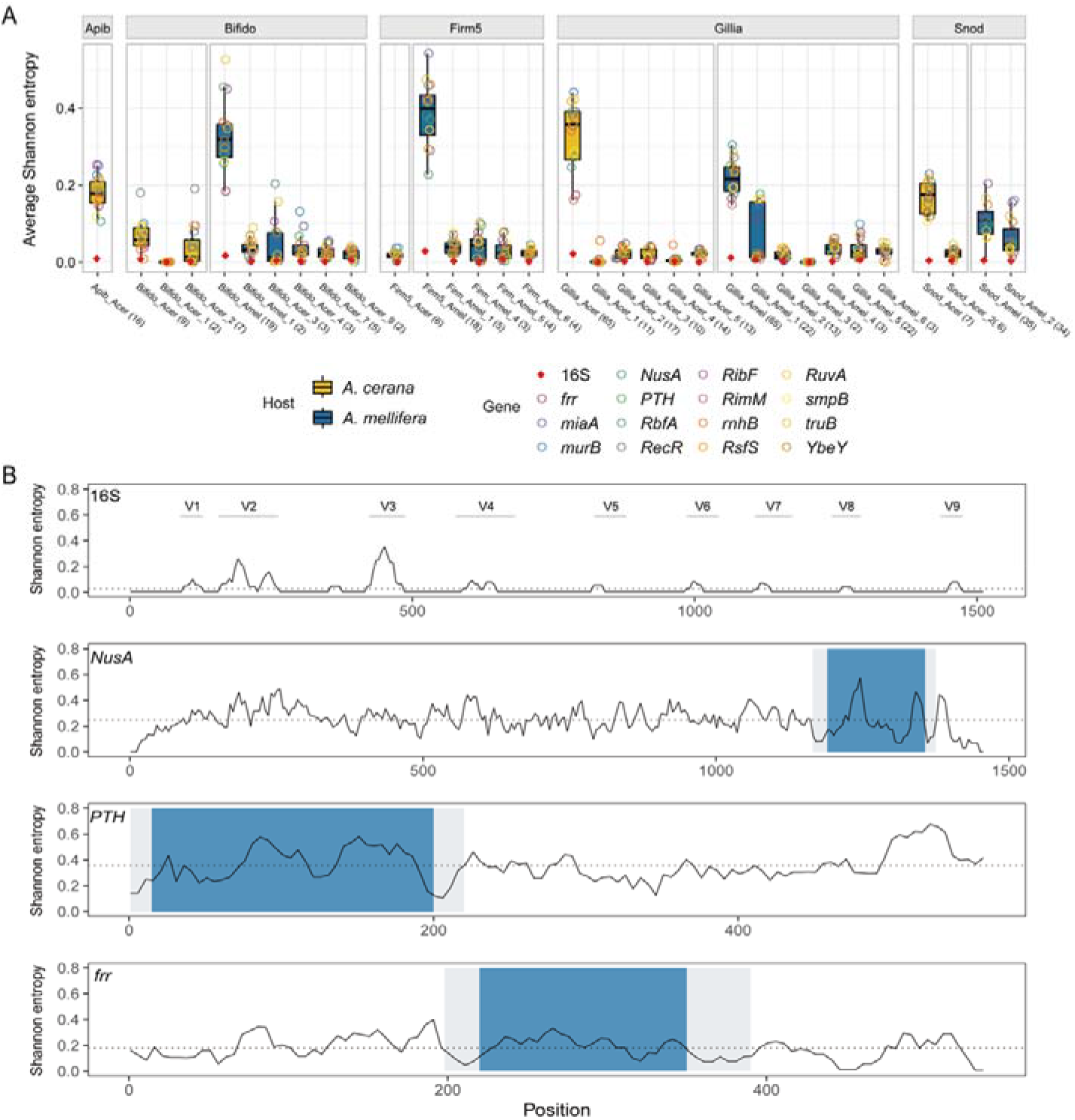
Marker genes are highly variable among SDPs. (A) Average Shannon entropy of the 15 marker genes and the 16S gene at both phylotype and SDP levels of honey bee gut bacteria. Numbers in brackets for each of the SDP groups indicate the number of strains examined for that specific group. (B) The Shannon entropy across 16S and candidate marker genes of all *A. cerana Gilliamella*. The Shannon entropy value is subsequently averaged by a 20-bp slide-window at a 5-bp step. Gray shadows depict conserved regions optimal for primer-binding sites and blue shadows are considered as hypervariable regions in this study. Dash lines represent the mean Shannon entropy values cross all sequences. Gray lines depict the classic variable regions of the 16S gene. Apib: *Apibacter*; Bifido: *Bifidobacterium*; Firm5: *Lactobacillus* Firm5; Gillia: *Gilliamella*; Snod: *Snodgrassella alvi*.

Because phylogenetic placement of the query sequence is a critical step in our SDP identification method, each marker gene will need to first produce a “correct” phylogeny for the phylotype in question. Therefore, we further examined whether each of the 15 marker genes could produce the same SDP phylogeny as inferred from whole-genome sequences of *Gilliamella*. Here, the tree based on all 65 *A. cerana Gilliamella* genomes was used as the gold standard. The results showed that all 15 marker genes but *rnhB* reconstructed the SDP phylogeny, with all strains assigned to corresponding SDPs (Fig. S2). On the *rnhB* gene tree, two *Gilliamella* genomes were misplaced from SDP Acer_Gillia_4 to Acer_Gillia_2, which was likely due to a higher sequence similarity between these two SDPs at a value of 90.93% ± 0.18 SD comparing to that between other SDPs (79.98% ± 1.89 SD). Therefore, *rnhB* was subsequently excluded from further screening.

For the 14 remaining marker genes, we further explored for regions that were suitable for amplicon sequencing, based on the presence of conserved primer regions flanking the hyper-variable region. The *RimM* gene lacked hyper variable regions across the full gene length (Fig. S1), while some other genes (*murB*, *RecR*, *miaA*, *RbfA*, *RibF*, *RuvA, RsfS* and *YebY*) did not demonstrate promising conserved regions for primer design. These genes were then excluded from the candidate gene pool. The 5 remaining candidates (*frr*, *NusA*, *PTH, truB* and *smpB*) all had a hyper-variable region of ~200-550 bp that was flanked by conservative primer regions. Among them, *frr, NusA* and *PTH* produced an amplicon of ~200 bp (Fig. 2B), which could be thoroughly sequenced with most current shotgun sequencing methods (e.g., PE100 or PE150). These 3 genes were then chosen for the final test for their performance in SDP discrimination in both identity and quantity, using *Gilliamella* mock samples and real honeybee guts.

### Marker gene amplicon sequencing (MGAS) showed high accuracy, sensitivity and repeatability in SDP profiling of mock samples

Mock samples contained varied proportions of the representative strain cultures of the 5 *Gilliamella* SDPs. These samples were extracted for DNA and amplified for the hyper-variable regions of the 3 candidate marker genes (*frr*, *NusA* and *PTH*). Twenty-four barcoded amplicons were pooled and shotgun sequenced for ca. 1 Gb data (ca. 2.5 million reads). Each mock sample was sequenced three times. An average of 73,462, 86,467 and 113,498 reads per sample was generated for *frr, NusA* and *PTH*, respectively.

The results of MGAS showed a high level of repeatability across the three replicates, where the average ICC(C,1) > 0.9, except for *PTH*, which had an ICC(C,1) of 0.752 among samples with equal proportion of bacterial DNA (Fig. 3A; Fig. S4C). With regards to detection accuracy, MGAS correctly detected all bacterial members present in 22/24 samples, while two samples (S03 and S04) showed false positive results, which was probably derived from sample contamination or sequencing error (Fig. 3B). Because the sensitivity of amplicon sequencing was affected by sequencing depth, we calculated the minimum read numbers required to detect members at low abundances, using rarefaction curves (Fig. S5). The results suggested that strains with a relative abundance of 1% could be detected by a minimum of ca. 1,123, 2,953 and 5,034 reads for *frr, NusA* and *PTH* (equivalent to 0.49, 1.29 and 2.44 Mb data per sample), respectively. Accordingly, lower abundance would require deeper sequencing. At a relative abundance of 0.02%, approximately 17,778, 18,518 and 22,222 reads (7.75, 8.07 and 10.76 Mb data) were required for *frr, NusA* and *PTH*, respectively (Fig. 3D; Fig. S5). The sequencing depth was generally sufficient for SDP detection in our study. Among the 216 sequenced samples, only two samples were sequenced with only 963 (*frr*) and 2,348 (*PTH*) reads, respectively, and failed in identifying corresponding SDP members at the lowest proportions (1% and 0.1%, respectively) due to insufficient sequencing depth.

**FIG 3.**
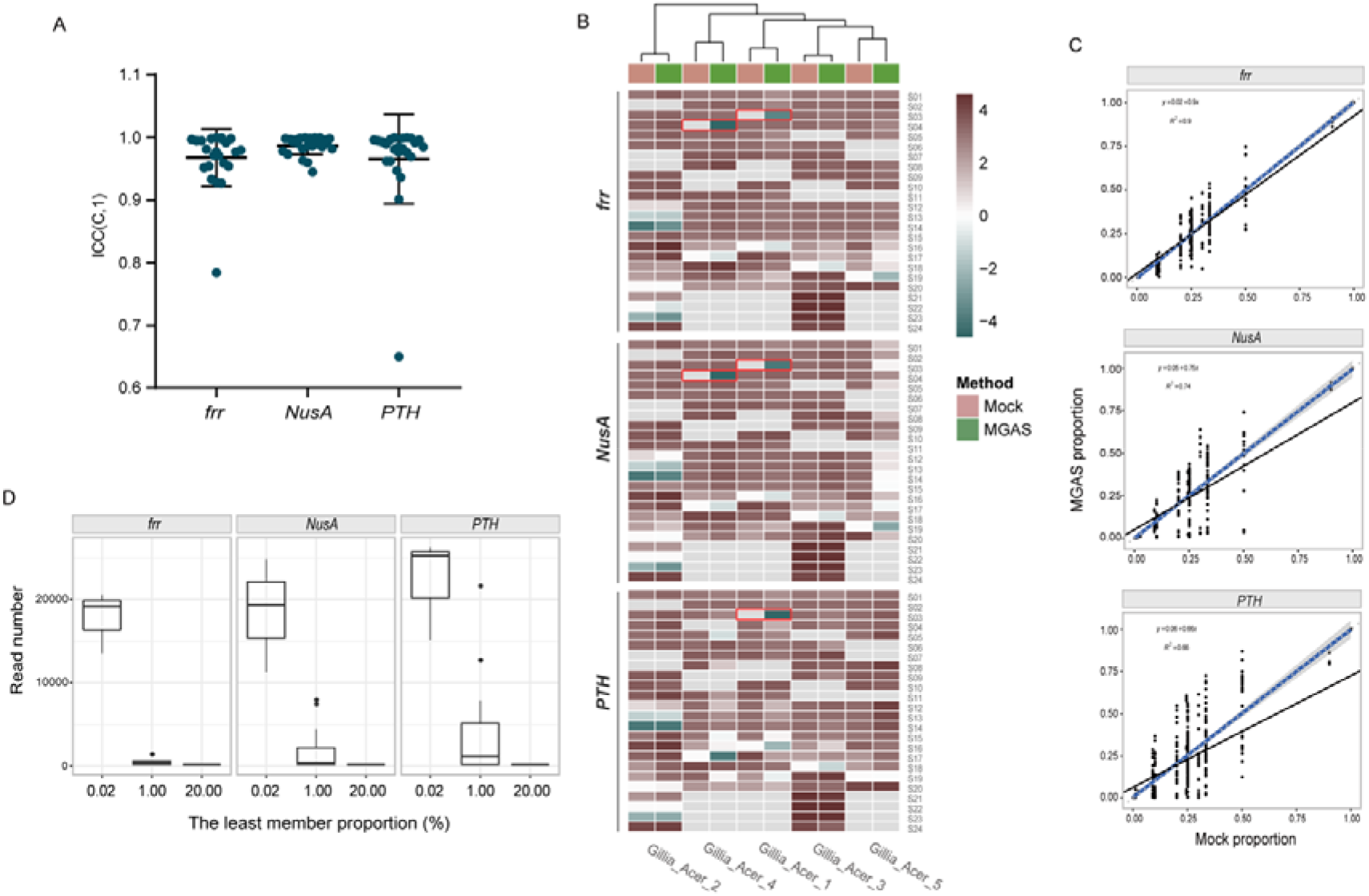
MGAS accurately identifies *A. cerana Gilliamella* SDPs. (A) Intraclass correlation coefficient (ICC) of relative abundance among the three replicates of MGAS samples. The ICC is calculated using the two-way mixed effects model with consistency (C) as the relationship among replicates, and single (1) result as the unit of measurement, i.e., ICC(C, 1). (B) Relative SDP abundances in mock samples revealed by marker gene sequencing. The results shown in the heatmap are the logarithms of the relative abundances of the five representative strains of the five SDPs of *A. cerana Gilliamella*. Grey box indicates a relative abundance at zero. False positive results are framed in red. (C) Spearman correlation of SDP abundances in *A. cerana Gilliamella* communities revealed by sequencing against mock samples. *p* <2.2e-16. The black line presents the linear regression of the MGAS results against SDP abundances in mock samples. The blue solid and gray dashed lines represent a 1: 1 line and the fitted exponential regression (with 95 % confidence interval shown in gray shade), respectively. (D) Minimum read numbers required for detecting members at low abundances.

In addition to accurately identify *Gilliamella* SDPs, all three marker genes performed well in quantifying relative abundances for mock samples. The relative abundances revealed by amplicon reads were highly congruent with corresponding mock proportions in bacterial mock samples, with the average R^2^ values of 0.91, 0.74 and 0.66 for *frr, NusA* and *PTH*, respectively *(p* < 2.2e-16, Fig. 3C). The DNA mock samples yielded similar results, with the average R^2^ values of 0.99, 0.91 and 0.99, for *frr, NusA* and *PTH*, respectively (Fig. S4B). Taken together, the MGAS method showed high levels of accuracy, sensitivity and repeatability in characterizing SDP compositions, in both taxonomic identity and relative abundance.

### MGAS performed equally well as metagenomics in characterizing honeybee gut SDP diversity

To examine the performance of the MGAS method in characterizing honeybee gut microbiota, we used *frr* (Fig. 4) and *PTH* (Fig. S6) genes to calculate *Gilliamella* SDP diversities for the 12 *A. cerana* workers from Sichuan and Taiwan, China. The MGAS was able to assign strains to the correct SDP at accurate abundance for real gut samples, with results were highly congruent with those from metagenomic sequencing (with R^2^ = 0.99 for *frr* and 0.97 for *PTH, p* < 2.2e-16, Fig. 4B; Fig. S6B). Both results revealed that most individual bees were dominated by two or three *Gilliamella* SDPs, yet with significant variations in dominant members and compositions among individuals and across geographical locations (Fig. 4A). Gillia_Acer_2 was the dominant SDP in most of the sequenced bees, which was found in 11 out of the 12 samples, with 10 bearing relative abundances of 48.06 - 98.37% (Fig. 4A). Both methods showed congruent results in alpha diversity (*p* = 0.82 and 0.79 for MGAS and metagenomics sequencing, respectively, Wilcoxon rank-sum test, Fig. 4C). At the beta diversity level, the principal coordinate analysis (PCoA) based on Bray-Curtis dissimilarity revealed that the gut bacterial communities from bees of Sichuan and Taiwan formed two distinct clusters, which separated along the first axis (Fig. 4D). This result was again consistent between the MGAS and metagenomic methods (Adonis PERMANOVA, R^2^ = 0.056, *p* = 0.204 for MGAS and R^2^ = 0.096, *p* = 0.134 for metagenomics). Thus, the performance of SDP profiling using MGAS was parallel to the metagenomic gold standard in microbial community studies.

**FIG 4.**
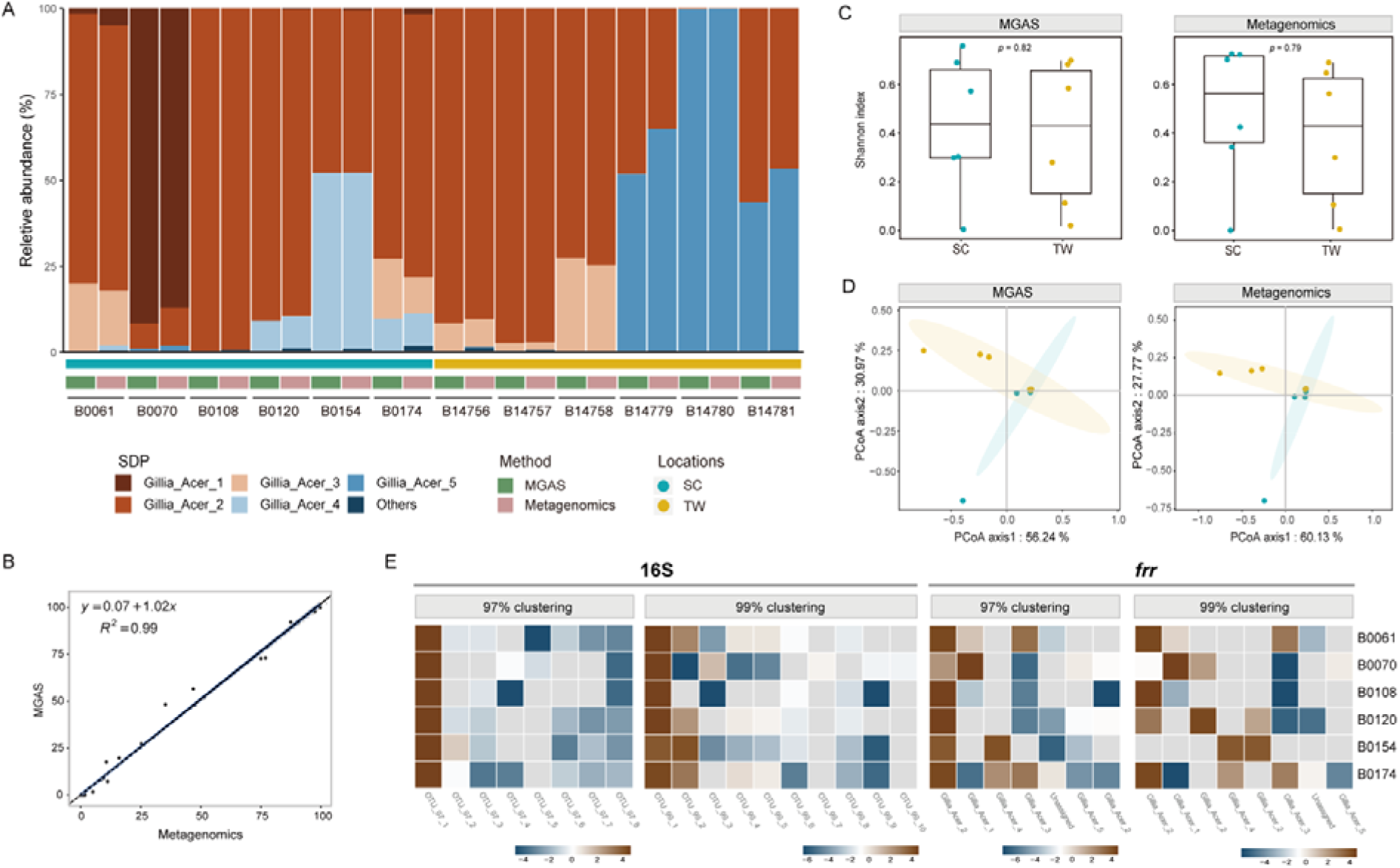
MGAS shows high congruence to metagenomic sequencing at SDP-level analysis. (A) Relative abundances of *Gilliamella* SDPs revealed by MGAS (*frr*) and metagenomics sequencing of *A. cerana* gut communities. (B) Spearman correlation coefficient between MGAS and metagenomics results, with R^2^ = 0.99, *p* < 2.2e-16. The black line presents the linear regression of the MGAS results in SDP abundances against those of metagenomics. The blue solid and gray dashed lines represent a 1: 1 line and the fitted exponential regression (with 95 % confidence interval shown in gray shade), respectively. (C) Shannon diversity index of SDP frequencies for bee guts from two locations calculated by MGAS (left panel) and metagenomic sequencing (right panel). The two methods showed no significant difference, with the *p*-value of 0.70 and 0.82 in SC and TW, respectively, by Wilcoxon rank-sum test. (D) Principal coordinate analysis (PCoA) based on Bray-Curtis dissimilarity of SDP compositions of honey bee workers from Sichuan and Taiwan using MGAS (left panel, Adonis PERMANOVA, R^2^ = 0.056, *p* = 0.204) and metagenomic sequencing (right panel, Adonis PERMANOVA, R^2^ = 0.096, *p* = 0.134). Each point represents the value for an individual bee and the color represent the location (Sichuan or Taiwan) of each bee. The shaded ellipses represent 95% confidence intervals on the ordination. (E) Relative abundances of *Gilliamella* OTUs in the gut microbiota of *A. cerana* assigned by clustering at 97% or 99% thresholds for 16S V4 and *frr*. The result shown in the heatmap are the logarithms of the relative abundances of the OTUs or five SDPs. Individual bees are marked to right of each row. Grey box indicates a relative abundance at zero.

The 16S V4 region was also used to determine the *Gilliamella* SDP compositions for the 6 bee gut samples from Sichuan. We applied operational taxonomic unit (OTU) clustering based on sequence similarity at 97% and 99% identity thresholds, which are commonly adopted for surveying phylotype and intra-phylotype microbial diversities, respectively [12, 29], to assess the efficacy of 16S in SDP profiling. 16S amplicon sequencing resulted in 8 and 10 OTUs at 97% and 99% thresholds, respectively, with a frequency cut off at > 100. The identified OTU numbers differed from those of the MGAS results at the same sequence similarity thresholds (Fig. 4E). Alarmingly, 16S amplicons failed to assign OTUs to the correct SDPs via blast. And the relative OTU proportions revealed by 16S disagreed with those from MGAS, where the numbers of dominant OTUs (> 1%) revealed by MGAS were more congruent to those from metagenomics. The improved performance with the MGAS method in characterizing SDP diversity is likely due to greater sequence divergence of the marker genes. For instance, the average pairwise inter-SDPs sequence similarity in the *frr* hyper-variable region was significantly lower (90.92% ± 3.18, n = 65) than that of the 16S rRNA gene V4 region (99.95% ± 0.65, n = 44) (Wilcoxon rank-sum test, *p* < 2e-16).

## SUMMARY AND DISCUSSION

We developed a pipeline to identify reliable marker genes for accurate identification and quantification of SDPs from bacterial communities. Three important criteria were applied in the assessment: the extent of sequence divergence, phylogenetic accuracy, and the presence of flanking conservative primer regions. Single-copy protein-coding genes identified by our pipeline were applied as marker genes in SDP quantification of honeybee gut microbiota, successfully producing results consistent with those from metagenomics, which were used as the gold standard. Conversely, we showed that the widely used 16S contained limited sequence divergence within phylotypes, failing to provide sufficient resolution in differentiating SDPs. As a result, 16S V4 amplicon sequencing cannot reflect fine scale bacterial diversity for the community. Consequently, dominant OTUs delineated by 16S at 97% or 99% thresholds significantly differed from the defined SDPs. On the other hand, the OTUs of single-copy protein-coding genes screened out by our pipeline were successfully assigned to the correct SDPs, and the numbers of dominant OTUs showed more congruent results to those from metagenomics.

Compared with whole-genome shotgun sequencing, amplicon sequencing of single-copy protein-coding genes provides an alternative solution to characterize SDP diversity in an accurate, quantitative and economical way. We address that not every single copy protein-coding gene is efficacious in SDP quantification. The candidate gene must meet all three criteria integrated in our pipeline to be a good marker gene. For a phylotype that is well represented by genomes of various lineages, all single-copy genes, including protein-coding genes, can be evaluated by our pipeline. In this case, we expect dozens to hundreds of proper marker genes to be filtered out. On the other hand, a small set of core single-copy protein-copy genes that are determined to be universally present among known bacteria, such as the 15 marker genes tested in this study, will likely provide candidate genes suitable for accurate characterization of SDP diversity for less known bacterial taxa.

Accurate identification of the SDP composition will also facilitate the prediction of the functional capacity of microbial communities. Functional attributes of a given bacterial lineage are strongly correlated to its phylogenetic position [30]. Therefore, various approaches, e.g., PICRUTs [31], have been developed to predict potential functions of a given microbial community based on phylogenetic profiles of bacterial members. However, 16S sequences are employed in most current programs for phylogenetic reconstruction. As demonstrated in this study, single-copy protein-coding genes identified by our pipeline show better fidelity in revealing phylogenetic relationships for the focal phylotype. Therefore, we anticipate that function prediction for microbial communities will be further improved by integrating single-copy protein-coding genes and the screening pipeline described here.

## MATERIALS AND METHODS

### Genome references of core gut bacteria of honeybees

A total of 242 bacterial genomes associated with *A. mellifera* and *A. cerana* were downloaded from the NCBI genome database (Table S1). These 242 genomes were used as the reference database of honeybee gut bacteria, which comprised the 6 major phylotypes: *Apibacter* (n=16), *Bifidobacterium* (n=28), *Lactobacillus* Firm4 (n=2), *Lactobacillus* Firm5 (n=24), *Gilliamella* (n=130) and *Snodgrassella* (n=42).

### SDP delineation for honeybee core phylotypes

Protein-coding genes of all sequenced genomes were annotated using Prokka (https://github.com/tseemann/prokka) [32]. Core genes, which were defined as being shared by > 99% strains of a given phylotype, were identified using Roary (version 3.13.0) [33] with the parameter -blastp 75. Multiple sequence alignments were carried out using MAFFT (version v7.467, https://github.com/The-Bioinformatics-Group/Albiorix/wiki/mafft) [34]. Phylogenetic trees were constructed using core single-copy genes of each phylotype by RAxML (version 8.2.12, -x 12345 -N 1000 -p 12345 -f a -m GTRGAMMA) [35]. Phylogenies were visualized in R (version 3.6.0) using the package ggtree_v2.4.1 [36] or iTOL (version 6.1.1) [37]. Pairwise genome-wide average nucleotide identity (gANI) values were calculated using pyani (version 0.2.10; https://github.com/widdowquinn/pyani) [38]. A clade with a gANI ≥ 95% from its closest clade was defined as an SDP.

### Screening for candidate marker genes capable of discriminating *Gilliamella* SDPs

The fifteen universal single-copy maker genes (*frr*, *NusA*, *PTH*, *RbfA*, *RecR*, *rnhB*, *RibF*, *RimM*, *RsfS*, *RuvA, smpB, truB, miaA, murB* and *YebY*, listed in Table S2) [24] were evaluated as candidate genes. The sequences of candidate marker genes were retrieved by MIDAS (version 1.3.2) [24], whereas the 16S genes were retrieved from the reference genomes using an in-house script. The average Shannon entropy (ASE) of the full gene length was used to assess sequence variation between strains of inter- and intra-SDPs for all phylotypes, where the Shannon entropy for each nucleotide site across genomes in comparison was calculated using oligotyping (version 2.1) [39].

The phylotype *Gilliamella*, which contains the most genomes available for this study, was used as a proof of concept to examine the efficacy of marker genes in SDP differentiation. For each SDPs in phylotype *Gilliamella*, the Shannon entropy values were subsequently averaged for each 20-bp slide-window with a 5-bp step to evaluate the regional genetic divergence along the full length of the marker genes. Pairwise sequence similarities were determined by Clustal Omega [40].

From the candidate genes, potential marker genes that may efficiently distinguish all known SDPs of the *Gilliamella* phylotype were screened. The following criteria were followed: 1) the marker genes should contain conservative regions flanking the hyper-variable region for designing primers enabling recovery target phylotype; 2) the amplicon length is between ~150-550 bps; 3) the amplified region is sufficiently variable to allow the discrimination of SDPs; and 4) the primers are specific to the focal phylotype to avoid off-target amplifications. The aforementioned 15 marker genes were subject to these criteria, and 5 of them *(ffr, NusA, PTH, truB* and *smpB)* were selected as potential markers for identifying SDPs of *A. cerana Gilliamella*. Among these, three genes *(ffr, NusA* and *PTH*) were subjected to further testing as a proof of concept, because their amplicon lengths were 206, 206 and 230 bp, respectively, which were ideal for current shotgun sequencing platforms. To increase the throughput and cost efficiency, 24 amplicons were pooled for one sequencing run. The 5’ end of both forward and reverse primers were tagged with 6-bp unique barcode sequences (see Table S3) to distinguish positive and negative DNA strains, and to differentiate samples.

### Bacterial mock samples

One representative strain from each of the five *Gilliamella* SDPs associated with *A. cerana* was cultured at 35°C and 5% CO_2_ for 48 h, on heart infusion agar (HIA) medium containing 5% sheep’s blood [41]. To screen potential contaminations, the full-length 16S gene was amplified for each bacterial culture using universal primers 27F and 1492R [41] and was subject to Sanger sequencing. 16S sequences were checked against those of the reference strains for identification, before strains were mixed for mock samples. Each *Gilliamella* culture was adjusted to OD600 = 0.5. Twenty-four mock SDP communities were prepared by mixing up 2-5 of the representative strains at varied proportions. The compositions of the mock samples were set as: equal proportion of each of the five strains, equal proportion of four strains with the absence of one strain at a time, equal proportion of three strains with the absence of two randomly selected strains, and a series of varied compositions with relative abundances ranging from ca. 0.02% to 50%. DNA of the bacterial mixtures were extracted using a CTAB-based DNA extraction protocol followed by recovery in 10 mM Tris-EDTA buffer (1×TE, pH 7.4) and quantified using the Qubit^®^ DNA Assay Kit on a Qubit^®^ 3.0 Fluorometer (Life Technologies, CA, USA). Alternatively, genomic DNA of each of the five representative strain cultures was extracted separately and the mixed at varied compositions and proportions (see Table S4).

### SDP identification and quantification for mock samples using amplicon sequencing of the three marker genes

PCR amplification was performed for *frr* (frr-F 5’ GCTGAAGATGCAAGAAC and frr-R 5’ GCATCACGACGAATATT), *NusA* (NusA-F 5’ CTTGAAATTGAAGAACT and NusA-R 5’ GTACCTTGTTCAGCTAA), and *PTH* (PTH-F 5’ AAACTTATTGTAGG and PTH-R 5’ CCACTTAAATTCATAAA) for each mock sample with three replicates. Triplicate 50-μl reactions were carried out with 25 μl of 2 × Phanta Max Master Mix (Vazyme Biotech, Nanjing, China), 2 μl (each) of 10 μM primer, 19 μl of ddH_2_O, and 2 μl of template DNA. The thermocycling profile consisted of an initial 3-min denaturation at 95 °C, 35 cycles of 15 s at 95 °C, 15 s at 52 °C for *NusA* and *frr* or at 42 °C for *PTH*, and 20 s at 72 °C and a final 10-min extension step at 72 °C. After being visualized on 2% agarose gels, DNA was purified using a gel extraction kit (Qiagen, Germany) and quantified using the Qubit^®^ DNA Assay Kit on a Qubit^®^ 3.0 Fluorometer. Barcoded amplicons of up to 24 mock samples were pooled together and subject to Illumina sequencing using a NovaSeq 6000 platform (PCR-free library, 150 PE) at Novogene (Beijing, China). Approximately 1 Gb of raw data were obtained from each pooled library (Table S5).

The program fastq-multx (version 1.3.1. https://github.com/brwnj/fastq-multx) was employed to demultiplex sequencing reads based on barcode sequences. The 6-bp barcodes in reverse sequences were trimmed using Seqtk (https://github.com/lh3/seqtk). The demultiplexed paired-end reads were then analyzed in QIIME2 (version 2020.2. https://qiime2.org) [42]. A plugin DATA2 [43] was used to denoise reads and to group sequences into amplicon sequence variants (ASVs). Individual ASVs were then taxonomically classified using blast (classify-consensus-blast) at a 97% identity threshold (Fig. S3) against the 3 marker genes *(ffr, NusA* and *PTH)* derived from the customized bee gut bacterial dataset. The relative abundance of each SDP (RA_SDP_) was calculated as: RA_SDP_ = (NR_SDP_) / (NR_Gillia_)*100, where NR_SDP_ represents the number of reads mapped to the focal SDP and NR_Gillia_ represents the number of reads mapped to all *Gilliamella* SDPs. These estimated abundances were then compared to those of the mock samples. The performance of SDP profiling of the 3 marker genes was evaluated on the basis of accuracy, sensitivity and repeatability. Intraclass correlation coefficient (ICC) with a two way random/mixed (ICC(C,1)) model was used to assess the repeatability of this method using SPSS (version 20.1) [44].

Rarefaction curves were plotted using identified SDP numbers against read numbers, which were used to infer the minimum read number required to detect strains at varied proportions. For each sample, ASVs with a depth <100 were filtered out. Rarefaction was performed using QIIME2 with the plugin alpha-rarefaction and a sampling depth of 40,000 reads per sample and default parameters. Minimum read numbers for identifying SDPs with relative abundances of 0.02%, 1% and 20% were chosen manually.

### SDP identification and quantification for *A. cerana* gut microbiota using 16S, marker genes, and metagenome sequencing

Adult worker bees collected in Sichuan were used to quantify *Gilliamella* SDP diversity using three different methods (16S V4 region amplicon sequencing, MGAS and metagenomic sequencing). Bees were first cooled at 4 °C for 10 min. Then the entire guts were dissected from the abdomen using sterile forceps and DNA was extracted using a CTAB bead-beating protocol described previously [45].

Firstly, the 16S V4 region was amplified for six bee guts from Sichuan and sequenced using an Illumina Hiseq X Ten platform (250-300 bp insert size, 250 PE) at BGI-Shenzhen (Shenzhen, China). Raw reads obtained for each sample were summarized in Table S6. Data quality control was performed using fastp (version 0.13.1, -q 20 -u 10 -w 16) [46]. The demultiplexed sequences were denoised and grouped into ASVs using an open reference method VSEARCH [47] embedded in QIIME 2. The taxonomic identification for ASVs was subsequently performed using the naive-Bayesian classifier trained on the BGM-Db, a curated 16S reference database for the classification of honeybee and bumblebee gut bacteria [48]. A feature table and ASVs consisting of filtered 16S reads pertaining to *Gilliamella* was constructed. OTU clustering was performed at both 97% and 99% identity thresholds, respectively, using VSEARCH with cluster-features-de-novo method. Additionally, low-abundant OTUs comprising of <100 reads were removed. Taxonomic assignments for OTUs were performed using blast against the BGM-Db with SDP-level taxonomy. OTU composition heatmaps were generated based on relative abundances and visualized in R.

Secondly, for each sample, the marker genes *frr* and *PTH*, which demonstrated the best and worst performances in accuracy and sensitivity, respectively, among the 3 marker genes, were applied following the same pipeline used in the mock samples. ASVs of the six sample from Sichuan were clustered into OTUs and filtered following the abovementioned 16S V4 pipeline. Taxonomic assignments for OTUs were performed by blast against *frr* sequences derived from the customized bee gut bacterial genome sequence database.

Finally, metagenome sequencing of four bee (B0108, B0120, B0154 and B0174) guts was performed using an Illumina Hiseq X Ten platform (300-400 bp insert size, 150 PE) at BGI-Shenzhen. Additional metagenomes of eight worker bee guts (BioProject PRJNA705951) were download from NCBI (Table S6). The metagenome sequencing was used as the gold standard for *Gilliamella* diversity distributed in the honeybee guts. Shotgun reads mapped to the *A. cerana* genome (GCF_001442555.1) using BWA aln (version 0.7.16a-r1181, -n 1) [49] were identified as host reads and subsequently excluded. We used the ‘run_midas.py species’ script in MIDAS with default parameters to estimate the relative abundances of SDPs for each sample. Finally, the results from MGAS were compared to those from metagenome sequencing to assess the performance of the marker genes.

### Data availability

Raw data from MGAS, 16S V4 amplicon and metagenomic sequencing have been submitted to NCBI under BioProject PRJNA772085.

## SUPPLEMENTAL MATERIAL

## ACKNOWLEDGEMENTS

This work was supported by the Program of Ministry of Science and Technology of China (2018FY100403), National Natural Science Foundation of China (No. 31772493) and National Natural Science Foundation of China (No. 32000346).

The authors declare no competing financial interests.

Xin Z. and Xue Z. designed, organized and coordinated the study. C.Y. conducted the screening pipeline development, marker gene and 16S V4 amplicon sequencing analysis. Q.S. retrieved the sequences of the 16S and single-copy protein-coding genes, and conducted reference-based metagenome mapping. M.T. assisted in sequence variation analysis. S.L. conducted sample collection and SDP identification. Xin Z., Xue Z., C.Y. and Hao Z. wrote the first drafts and all authors contributed to and proofed the manuscript.

---

**FIG S1.**
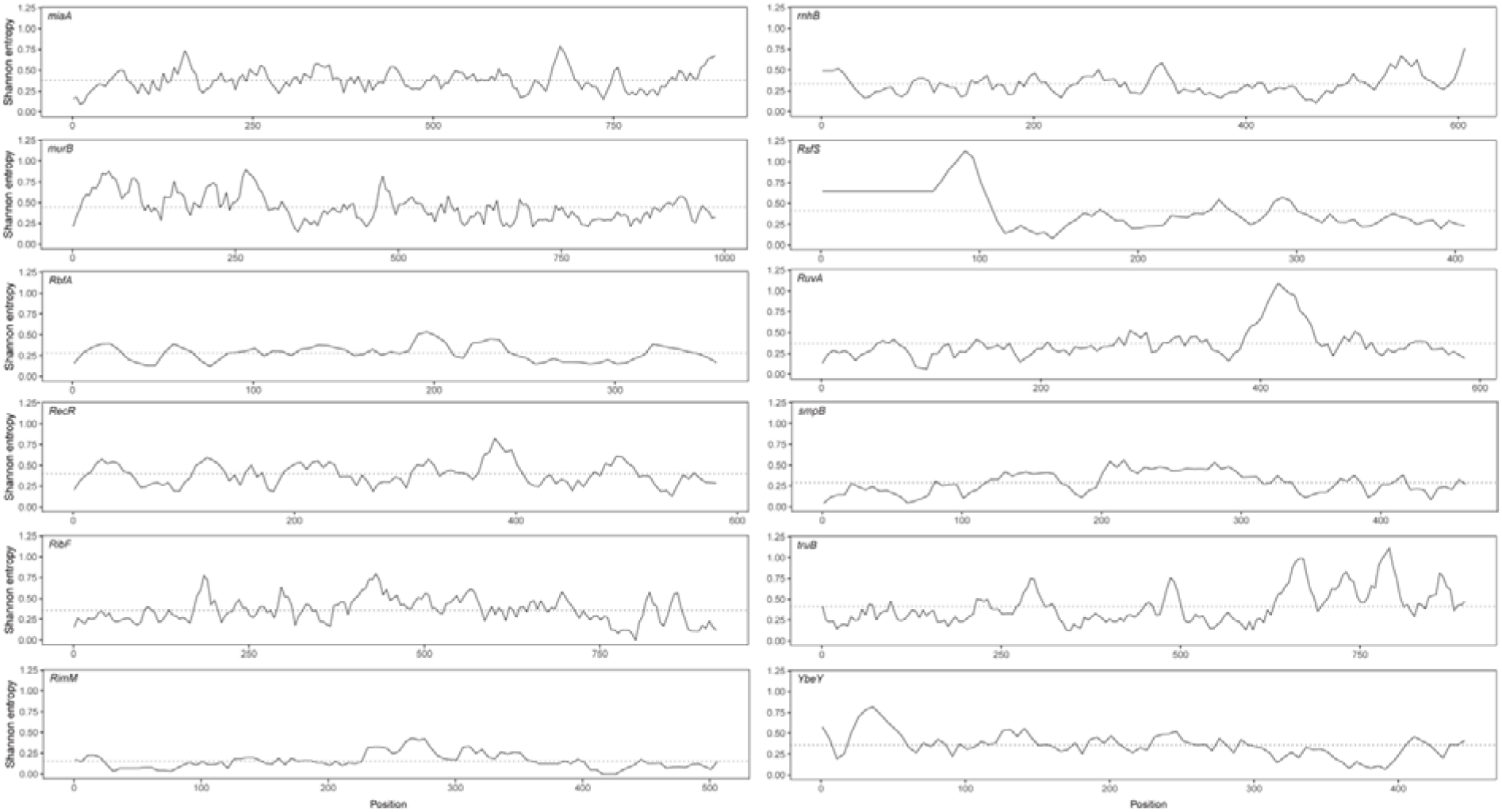
The Shannon entropy across the remain marker genes of all *A. cerana Gilliamella*. The Shannon entropy value is subsequently averaged by a 20-bp slide-window at a 5-bp step. Dash lines represent the mean Shannon entropy values cross all sequences.

**FIG S2.**
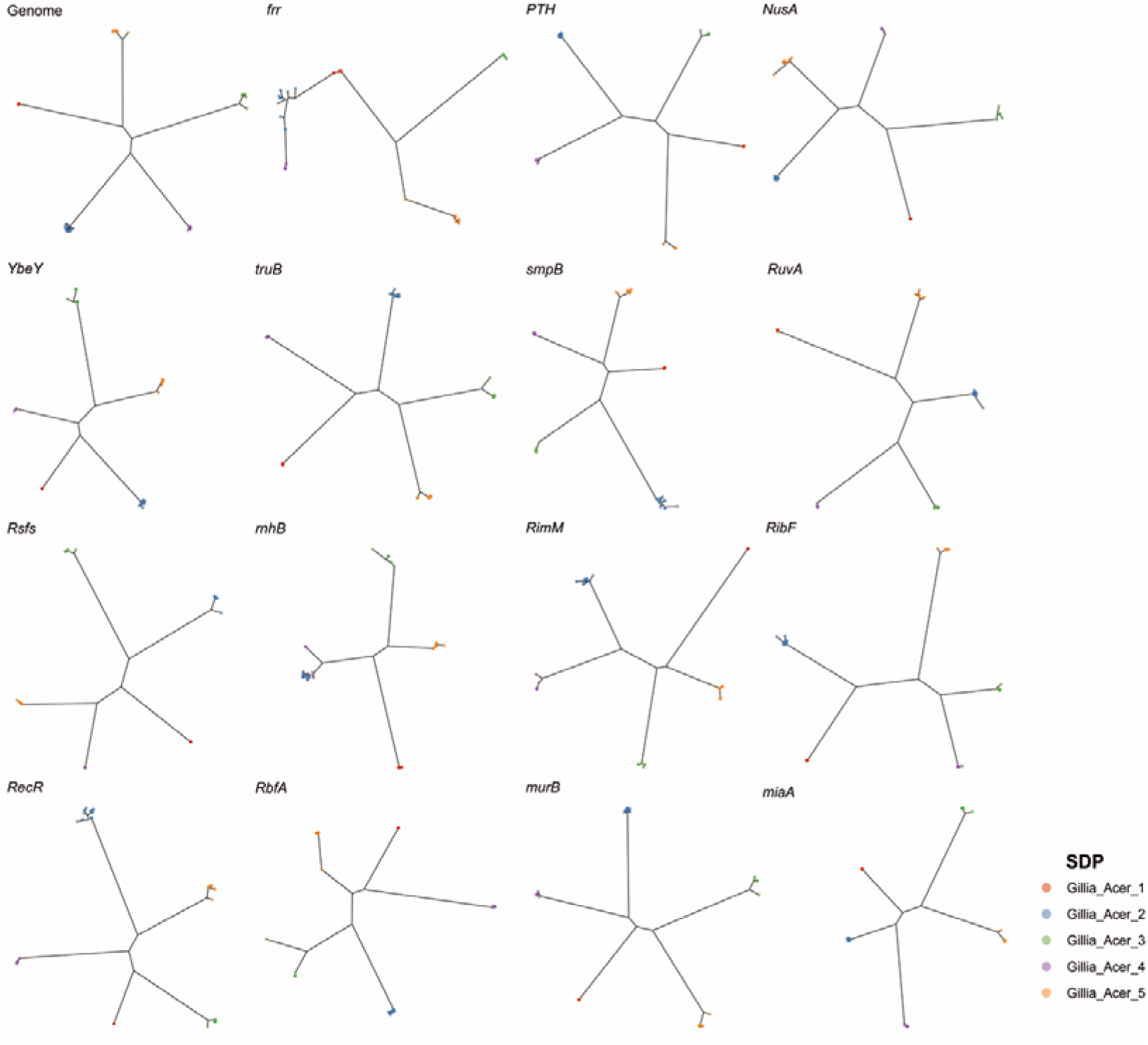
All but *rnhB* of the 15 marker genes produce five SDPs for *A. cerana Gilliamella* phylotype in concert with the whole-genome result.

**FIG S3.**
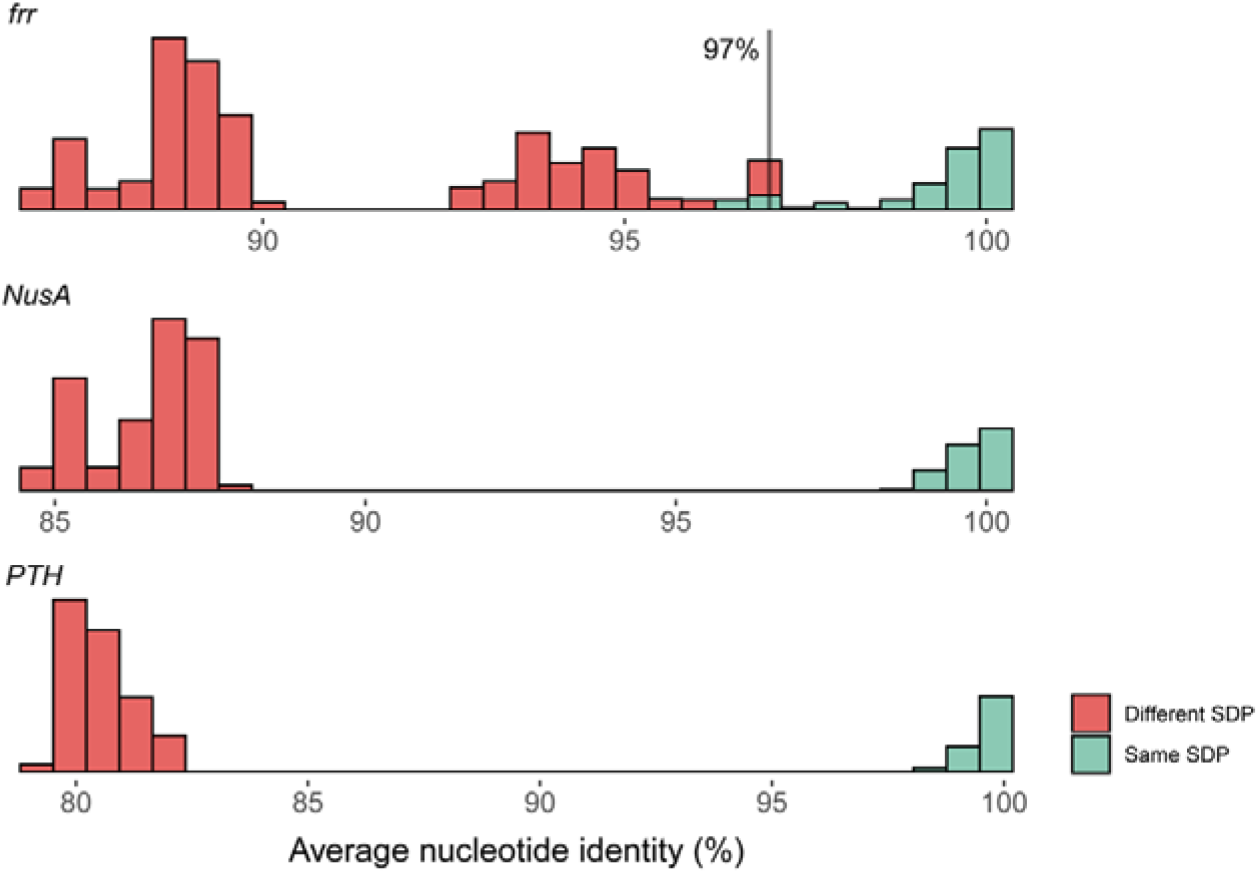
Histograms of average nucleotide identity values of the 3 marker genes from comparisons between strains belonging to the same SDPs (green) or different SDPs (red). Vertical black line indicates the threshold for bacterial SDPs taxonomy for the present method.

**FIG S4.**
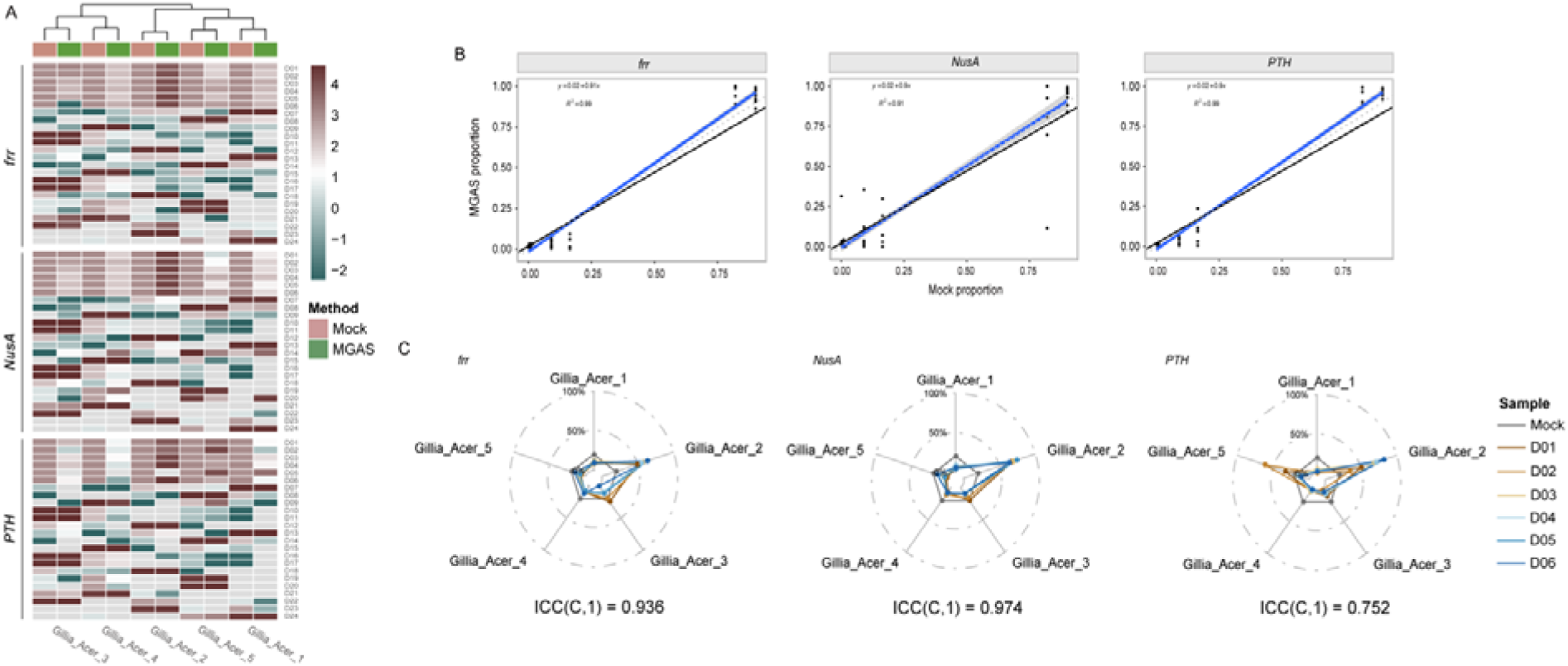
MGAS accurately identifies the *A. cerana Gilliamella* SDPs in DNA mock samples. (A) Relative SDP abundances in mock samples revealed by MGAS. The results shown in the heatmap are the logarithms of the relative abundances percentage of the five representative strains of the five SDPs of *A. cerana Gilliamella*. Grey box indicates a relative abundance at zero. (B) Spearman correlation of SDP abundances in *A. cerana Gillimella* communities revealed by sequencing against mock samples, *p* < 2.2e-16. The black line presents the linear regression of the MGAS results against SDP abundances in mock samples. The blue solid and gray dashed lines represent a 1: 1 line and the fitted exponential regression (with 95 % confidence interval shown in gray shade), respectively. (C) Repeatability of relative abundance between replicates of DNA mock samples. n = 6, ICC(C,1) is 0.936, 0.974 and 0.752 for *frr, NusA* and *PTH* genes, respectively.

**FIG S5.**
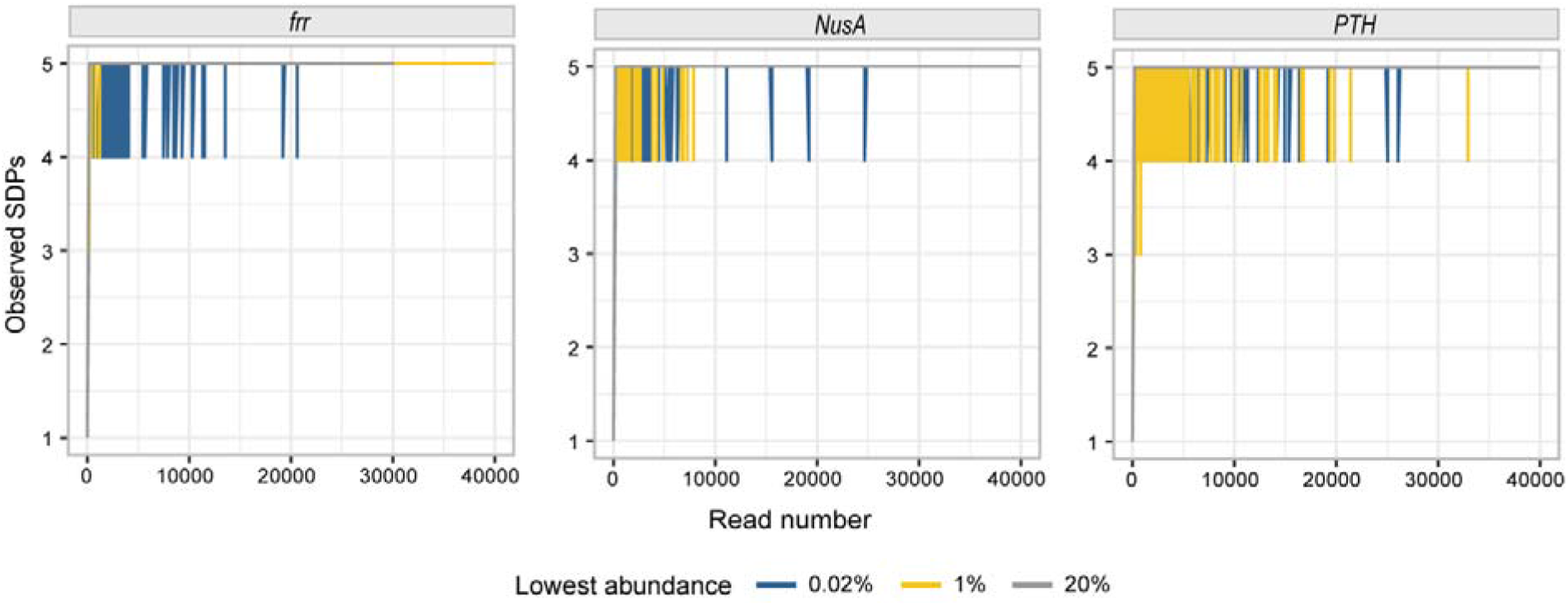
Rarefaction curves of detected bacterial SDPs in bacterial mock samples reach the saturation stage with increasing read numbers.

**FIG S6.**
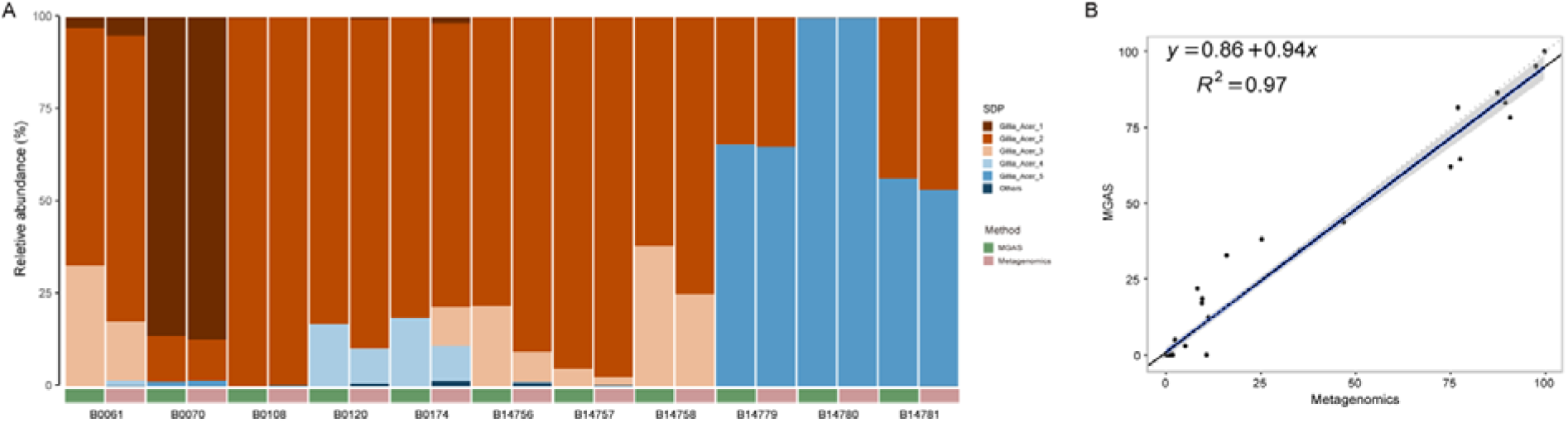
Amplicon sequencing with the *PTH* gene showed high congruence to metagenomic sequencing at SDP-level analyses. (A) Relative abundances of *Gilliamella* SDPs revealed by MGAS (*PTH* gene) and metagenomics sequencing of *A. cerana* gut communities. (B) Spearman correlation coefficient between MGAS and metagenomics results, with R^2^ = 0.97, *p* < 2.2e-16. The black line presents the linear regression of the MGAS results in SDP abundances against those of metagenomics. The blue solid and gray dashed lines represent a 1: 1 line and the fitted exponential regression (with 95 % confidence interval shown in gray shade), respectively.

**TABLE S1.**
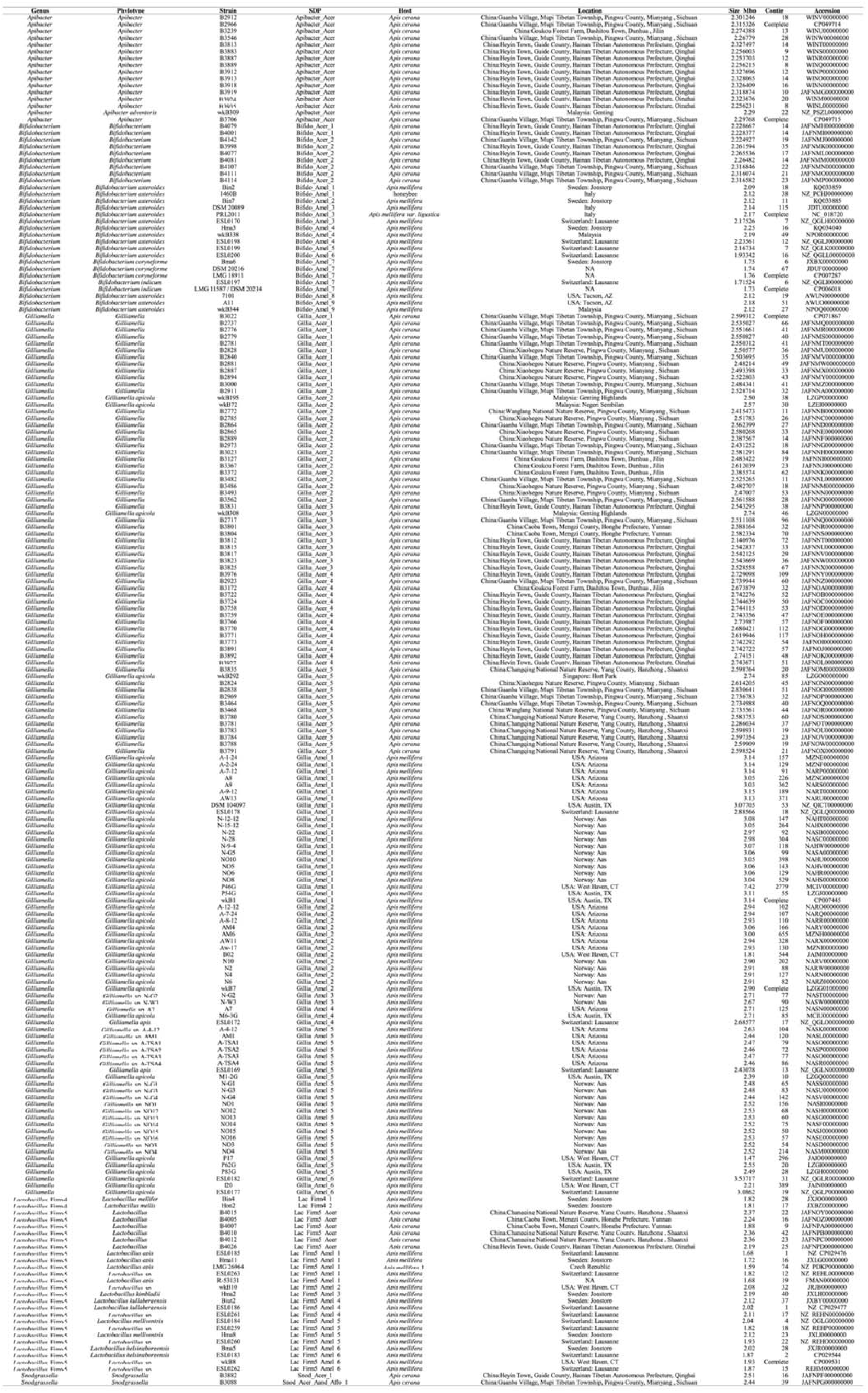
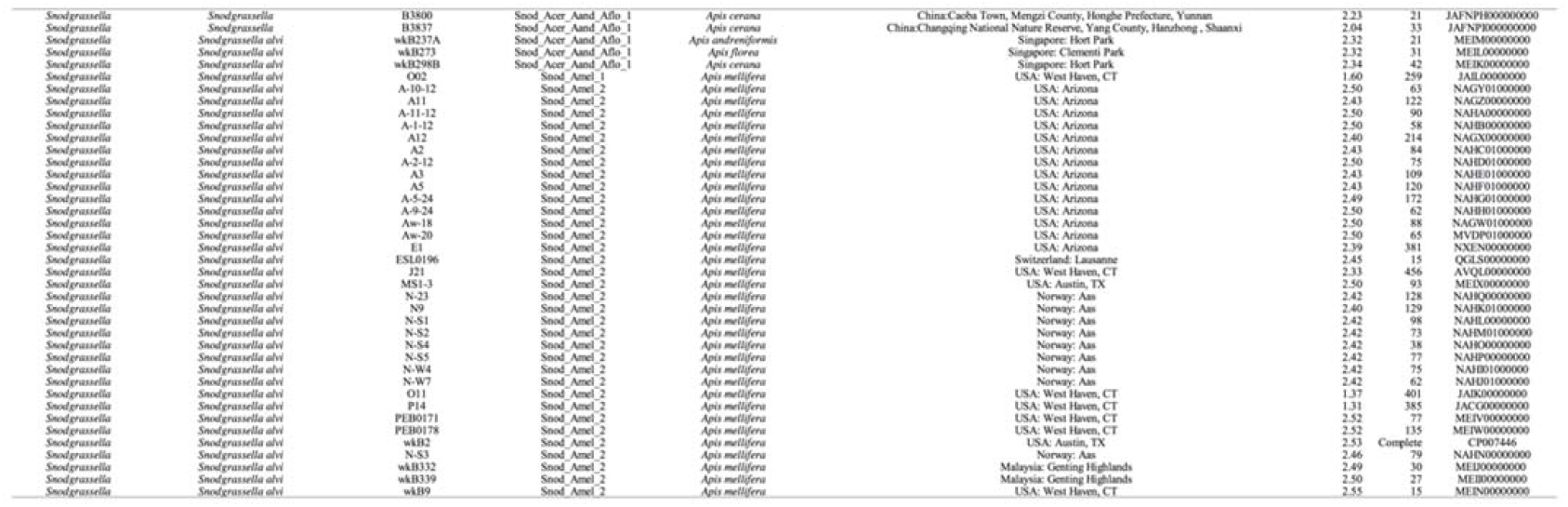
Information of the reference genomes.

**TABLE S2.**
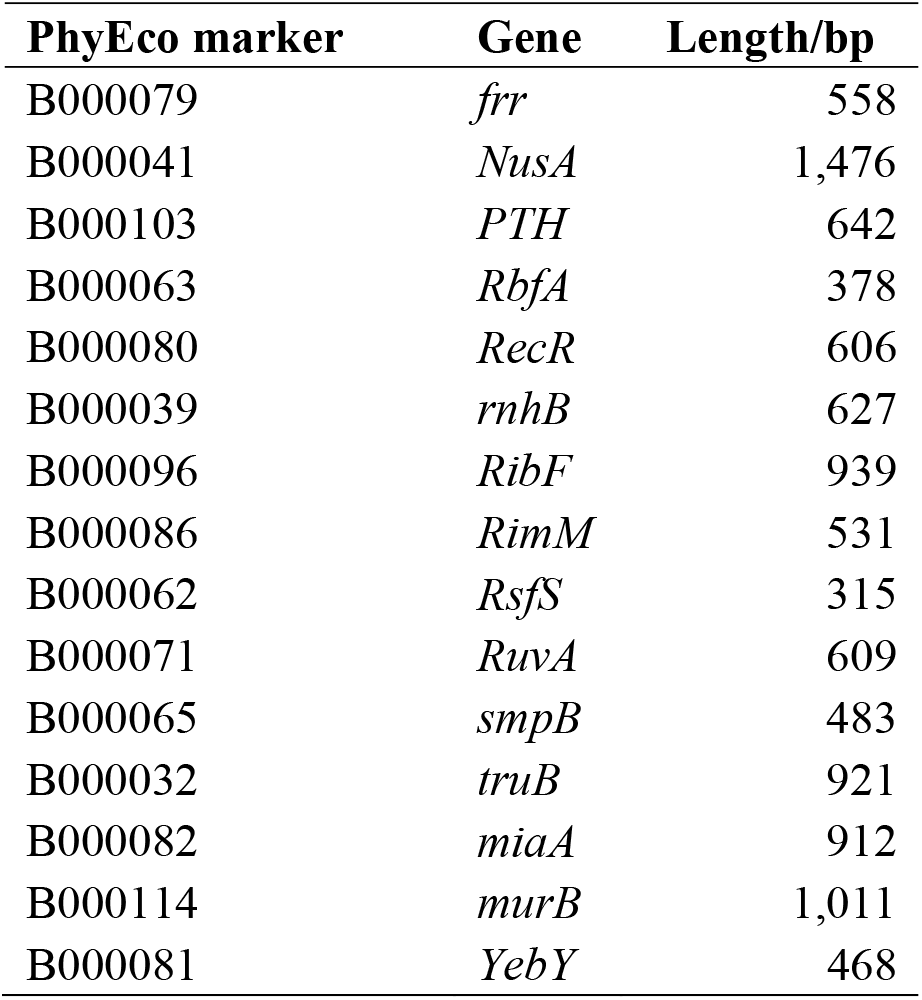
Information of the marker genes.

| **PhyEco marker** | **Gene** | **Length/bp** |
| --- | --- | --- |
| B000079 | *frr* | 558 |
| B000041 | *NusA* | 1,476 |
| B000103 | *PTH* | 642 |
| B000063 | *RbfA* | 378 |
| B000080 | *RecR* | 606 |
| B000039 | *rnhB* | 627 |
| B000096 | *RibF* | 939 |
| B000086 | *RimM* | 531 |
| B000062 | *RsfS* | 315 |
| B000071 | *RuvA* | 609 |
| B000065 | *smpB* | 483 |
| B000032 | *truB* | 921 |
| B000082 | *miaA* | 912 |
| B000114 | *murB* | 1,011 |
| B000081 | *YebY* | 468 |

**TABLE S3.** List of barcode sequences.

| **Barcode NO.** | **Forward seq (5'to 3')** | **Reverse seq (5'to 3')** |
| --- | --- | --- |
| B01 | ATCACG | ACTGAT |
| B02 | CGATGT | ATGAGC |
| B03 | TTAGGC | ATTCCT |
| B04 | TGACCA | CAAAAG |
| B05 | ACAGTG | CAACTA |
| B06 | GCCAAT | CACCGG |
| B07 | CAGATC | CACGAT |
| B08 | ACTTGA | CACTCA |
| B09 | GATCAG | CAGGCG |
| B10 | TAGCTT | CATGGC |
| B11 | GGCTAC | CATTTT |
| B12 | CTTGTA | CCAACA |
| B13 | AGTCAA | CGGAAT |
| B14 | AGTTCC | CTAGCT |
| B15 | ATGTCA | CTATAC |
| B16 | CCGTCC | CTCAGA |
| B17 | GTAGAG | GACGAC |
| B18 | GTCCGC | TAATCG |
| B19 | GTGAAA | TACAGC |
| B20 | GTGGCC | TATAAT |
| B21 | GTTTCG | TCATTC |
| B22 | CGTACG | TCCCGA |
| B23 | GAGTGG | TCGAAG |
| B24 | GGTAGC | TCGGCA |

**TABLE S4.**
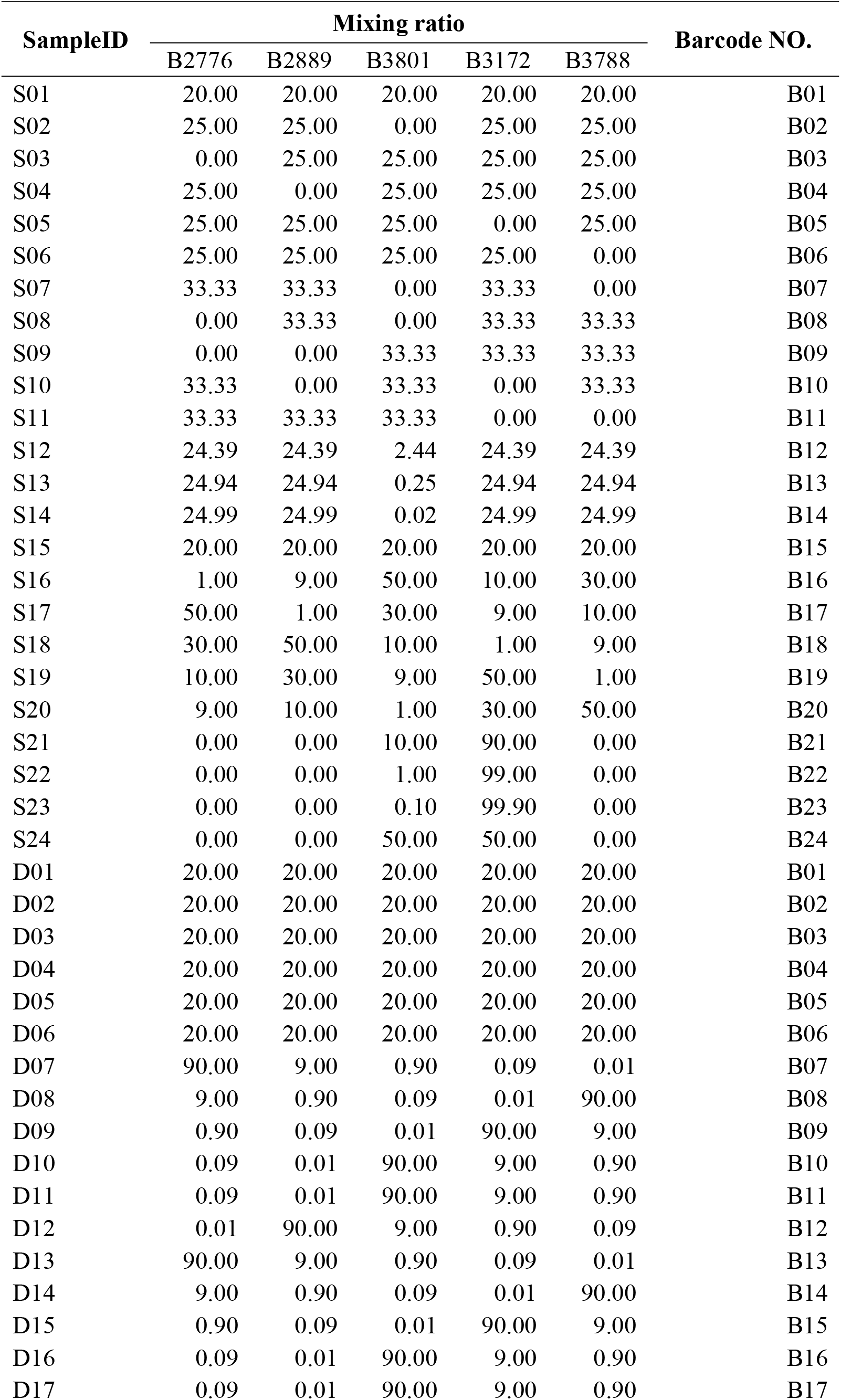
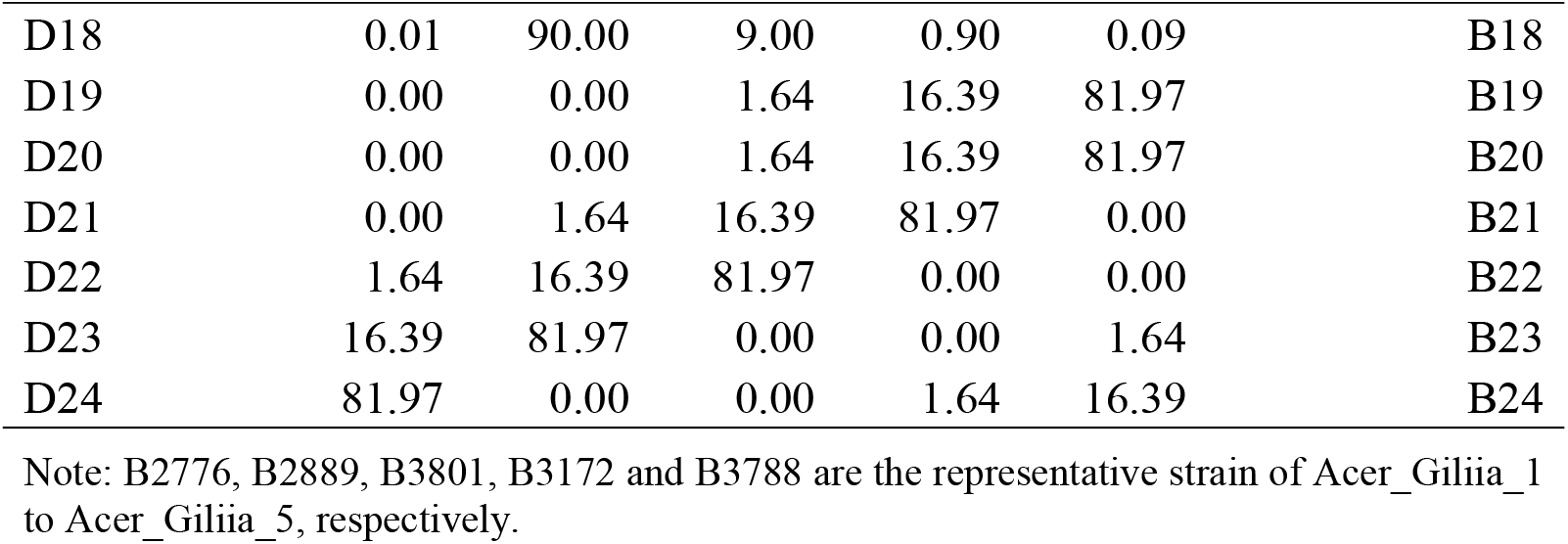
Mixing ratio of mock samples.

| SampleID | Mixing ratio |  |  |  |  | Barcode NO. |
| --- | --- | --- | --- | --- | --- | --- |
|  | B2776 | B2889 | B3801 | B3172 | B3788 |  |
| S01 | 20.00 | 20.00 | 20.00 | 20.00 | 20.00 | B01 |
| S02 | 25.00 | 25.00 | 0.00 | 25.00 | 25.00 | B02 |
| S03 | 0.00 | 25.00 | 25.00 | 25.00 | 25.00 | B03 |
| S04 | 25.00 | 0.00 | 25.00 | 25.00 | 25.00 | B04 |
| S05 | 25.00 | 25.00 | 25.00 | 0.00 | 25.00 | B05 |
| S06 | 25.00 | 25.00 | 25.00 | 25.00 | 0.00 | B06 |
| S07 | 33.33 | 33.33 | 0.00 | 33.33 | 0.00 | B07 |
| S08 | 0.00 | 33.33 | 0.00 | 33.33 | 33.33 | B08 |
| S09 | 0.00 | 0.00 | 33.33 | 33.33 | 33.33 | B09 |
| S10 | 33.33 | 0.00 | 33.33 | 0.00 | 33.33 | B10 |
| S11 | 33.33 | 33.33 | 33.33 | 0.00 | 0.00 | B11 |
| S12 | 24.39 | 24.39 | 2.44 | 24.39 | 24.39 | B12 |
| S13 | 24.94 | 24.94 | 0.25 | 24.94 | 24.94 | B13 |
| S14 | 24.99 | 24.99 | 0.02 | 24.99 | 24.99 | B14 |
| S15 | 20.00 | 20.00 | 20.00 | 20.00 | 20.00 | B15 |
| S16 | 1.00 | 9.00 | 50.00 | 10.00 | 30.00 | B16 |
| S17 | 50.00 | 1.00 | 30.00 | 9.00 | 10.00 | B17 |
| S18 | 30.00 | 50.00 | 10.00 | 1.00 | 9.00 | B18 |
| S19 | 10.00 | 30.00 | 9.00 | 50.00 | 1.00 | B19 |
| S20 | 9.00 | 10.00 | 1.00 | 30.00 | 50.00 | B20 |
| S21 | 0.00 | 0.00 | 10.00 | 90.00 | 0.00 | B21 |
| S22 | 0.00 | 0.00 | 1.00 | 99.00 | 0.00 | B22 |
| S23 | 0.00 | 0.00 | 0.10 | 99.90 | 0.00 | B23 |
| S24 | 0.00 | 0.00 | 50.00 | 50.00 | 0.00 | B24 |
| D01 | 20.00 | 20.00 | 20.00 | 20.00 | 20.00 | B01 |
| D02 | 20.00 | 20.00 | 20.00 | 20.00 | 20.00 | B02 |
| D03 | 20.00 | 20.00 | 20.00 | 20.00 | 20.00 | B03 |
| D04 | 20.00 | 20.00 | 20.00 | 20.00 | 20.00 | B04 |
| D05 | 20.00 | 20.00 | 20.00 | 20.00 | 20.00 | B05 |
| D06 | 20.00 | 20.00 | 20.00 | 20.00 | 20.00 | B06 |
| D07 | 90.00 | 9.00 | 0.90 | 0.09 | 0.01 | B07 |
| D08 | 9.00 | 0.90 | 0.09 | 0.01 | 90.00 | B08 |
| D09 | 0.90 | 0.09 | 0.01 | 90.00 | 9.00 | B09 |
| D10 | 0.09 | 0.01 | 90.00 | 9.00 | 0.90 | B10 |
| D11 | 0.09 | 0.01 | 90.00 | 9.00 | 0.90 | B11 |
| D12 | 0.01 | 90.00 | 9.00 | 0.90 | 0.09 | B12 |
| D13 | 90.00 | 9.00 | 0.90 | 0.09 | 0.01 | B13 |
| D14 | 9.00 | 0.90 | 0.09 | 0.01 | 90.00 | B14 |
| D15 | 0.90 | 0.09 | 0.01 | 90.00 | 9.00 | B15 |
| D16 | 0.09 | 0.01 | 90.00 | 9.00 | 0.90 | B16 |
| D17 | 0.09 | 0.01 | 90.00 | 9.00 | 0.90 | B17 |
| D18 | 0.01 | 90.00 | 9.00 | 0.90 | 0.09 | B18 |
| D19 | 0.00 | 0.00 | 1.64 | 16.39 | 81.97 | B19 |
| D20 | 0.00 | 0.00 | 1.64 | 16.39 | 81.97 | B20 |
| D21 | 0.00 | 1.64 | 16.39 | 81.97 | 0.00 | B21 |
| D22 | 1.64 | 16.39 | 81.97 | 0.00 | 0.00 | B22 |
| D23 | 16.39 | 81.97 | 0.00 | 0.00 | 1.64 | B23 |
| D24 | 81.97 | 0.00 | 0.00 | 1.64 | 16.39 | B24 |

Note: B2776, B2889, B3801, B3172 and B3788 are the representative strain of Acer\_Giliia\_1 to Acer\_Giliia\_5, respectively.

**TABLE S5.** Statistics of data outputs.

| LibraryID | Raw reads | Clean reads | Raw base/G | Clean base/G | Effective rate/% | Q20/% | Q30/% | GC content/% |
| --- | --- | --- | --- | --- | --- | --- | --- | --- |
| f1S01-f1S24 | 2,910,358 | 2,904,839 | 0.87 | 0.87 | 99.81 | 98.96 | 96.8 | 42.92 |
| f2S01-f2S24 | 4,370,025 | 4,362,026 | 1.31 | 1.31 | 99.82 | 98.99 | 96.88 | 42.85 |
| f3S01-f3S24 | 3,971,727 | 3,966,181 | 1.19 | 1.19 | 99.86 | 98.34 | 94.65 | 42.84 |
| N1S01-N1S24 | 3,101,708 | 3,097,334 | 0.93 | 0.93 | 99.86 | 98.83 | 96.33 | 38.67 |
| N2S01-N2S24 | 3,455,304 | 3,451,312 | 1.04 | 1.04 | 99.88 | 97.96 | 93.59 | 38.66 |
| N3S01-N3S24 | 2,893,355 | 2,889,594 | 0.87 | 0.87 | 99.87 | 97.8 | 93.25 | 38.64 |
| P1S01-P1S24 | 5,446,708 | 5,439,697 | 1.63 | 1.63 | 99.87 | 99.1 | 96.48 | 36.74 |
| P2S01-P2S24 | 2,698,030 | 2,694,490 | 0.81 | 0.81 | 99.87 | 98.97 | 96.29 | 36.88 |
| P3S01-P3S24 | 3,377,356 | 3,371,139 | 1.01 | 1.01 | 99.82 | 97.95 | 93.16 | 37.08 |
| fD01-fD24 | 3,599,515 | 3,595,846 | 1.08 | 1.08 | 99.9 | 98.41 | 95.28 | 42.44 |
| ND01-ND24 | 4,399,592 | 4,393,529 | 1.32 | 1.32 | 99.86 | 98.42 | 95.02 | 38.73 |
| PD01-PD24 | 3,387,737 | 3,380,942 | 1.02 | 1.01 | 99.8 | 98.53 | 95.13 | 36.64 |
| fB0061-fB14781 | 5,441,182 | 5,434,806 | 1.63 | 1.63 | 99.88 | 99.24 | 97.36 | 42.25 |
| PB0061-PB14781 | 3,591,409 | 3,587,252 | 1.08 | 1.08 | 99.88 | 98.8 | 95.75 | 36.55 |

Note: 1. f1 - f3, N1 - N3 and P1 - P3 represent the three replicates for *frr*, *NusA* and *PTH* gene sequencing, and S01 - S24 are mock samples with different ratio of mixing bacterial cells shown in Table S4.
3. f and P represent the *frr* and *PTH* gene sequencing, and BXXXX present *Apis cerana* gut sample

**TABLE S6.** Summary of read processing and data obtained from marker gene, 16S V4 amplicon and metagenomic sequencing of honey bee guts.

| Gut ID | Raw PE reads |  |  |  | Joined and filtered reads |  |  |  | *Gilliamella* reads |  |
| --- | --- | --- | --- | --- | --- | --- | --- | --- | --- | --- |
|  | 16S | *frr* | *PTH* | Meta | 16S | *frr* | *PTH* | Meta | 16S | Meta |
| B0061 | 84,584 | 358,851 | 65,424 | 33,687,518 | 82,760 | 311,842 | 60,964 | 5,159,739 | 25,700 | 3,185 |
| B0070 | 85,169 | 253,390 | 94,393 | 37,685,288 | 83,491 | 223,179 | 91,436 | 5,909,974 | 47,892 | 7,661 |
| B0108 | 83,908 | 349,570 | 123,368 | 45,115,102 | 82,132 | 344,248 | 121,018 | 4,242,595 | 39,424 | 2,189 |
| B0120 | 85,368 | 389,432 | 29,023 | 34,543,934 | 83,267 | 262,273 | 28,277 | 4,523,366 | 22,741 | 815 |
| B0154 | 83,691 | 289,878 | - | 34,969,488 | 81,143 | 226,310 | - | 4,622,933 | 26,346 | 1,086 |
| B0174 | 84,281 | 361,728 | 75,882 | 38,471,836 | 81,979 | 287,357 | 68,222 | 6,058,633 | 18,258 | 243 |
| B14756 | - | 224,748 | 118,334 | 21,491,068 | - | 194,850 | 114,236 | 4,622,933 | - | 17,472 |
| B14757 | - | 354,956 | 158,879 | 22,959,909 | - | 328,188 | 156,087 | 4,622,933 | - | 5,546 |
| B14758 | - | 277,823 | 165,638 | 24,408,709 | - | 224,100 | 160,741 | 9,658,926 | - | 4,182 |
| B14779 | - | 342,928 | 151,481 | 23,802,654 | - | 285,197 | 146,421 | 7,922,065 | - | 8,133 |
| B14780 | - | 301,064 | 48,088 | 23,495,452 | - | 272,069 | 47,247 | 7,123,654 | - | 3,064 |
| B14781 | - | 291,415 | 71,187 | 22,381,871 | - | 237,037 | 68,190 | 9,626,197 | - | 3,686 |

Note: “16S” indicates 16S V4; “Meta” indicates metagenomic; “-” indicates no test.

## REFERENCES

1. Moreira D, López-García P. 2011. Phylotype, p 1254–1254. In Gargaud M, Amils R, Quintanilla JC, Cleaves HJ, Irvine WM, Pinti DL, Viso M. Berlin, Heidelberg, Encyclopedia of Astrobiology. Springer Berlin Heidelberg.

2. von Mentzer A, Connor TR, Wieler LH, Semmler T, Iguchi A, Thomson NR, Rasko DA, Joffre E, Corander J, Pickard D, Wiklund G, Svennerholm A-M, Sjöling Å, Dougan G. 2014. Identification of enterotoxigenic *Escherichia coli* (ETEC) clades with long-term global distribution. Nat Genet 46:1321–1326.

3. Jain C, Rodriguez RL, Phillippy AM, Konstantinidis KT, Aluru S. 2018. High throughput ANI analysis of 90K prokaryotic genomes reveals clear species boundaries. Nat Commun 9(1):5114.

4. Caro-Quintero A, Konstantinidis KT. 2012. Bacterial species may exist, metagenomics reveal. Environ Microbiol 14:347–355.

5. Ellegaard KM, Engel P. 2019. Genomic diversity landscape of the honey bee gut microbiota. Nat Commun 10:446.

6. Konstantinidis KT, Tiedje JM. 2005. Genomic insights that advance the species definition for prokaryotes. Proc Natl Acad Sci U S A 102:2567–2572.

7. Fehlner-Peach H, Magnabosco C, Raghavan V, Scher JU, Tett A, Cox LM, Gottsegen C, Watters A, Wiltshire-Gordon JD, Segata N, Bonneau R, Littman DR. 2019. Distinct polysaccharide utilization profiles of human intestinal *Prevotella copri* isolates. Cell Host Microbe 26:680–690 e685.

8. Tett A, Huang KD, Asnicar F, Fehlner-Peach H, Pasolli E, Karcher N, Armanini F, Manghi P, Bonham K, Zolfo M, Filippis FD, Magnabosco C, Bonneau R, Lusingu J, Amuasi J, Reinhard K, Rattei T, Boulund F, Engstrand L, Zink A, Collado MC, Littman DR, Eibach D, Ercolini D, Rota-Stabelli O, Huttenhower C, Maixner F, and Segata N. 2019. The *Prevotella copri* complex comprises four distinct clades underrepresented in westernized populations. Cell Host Microbe 26:666–679 e667.

9. Oh S, Caro-Quintero A, Tsementzi D, DeLeon-Rodriguez N, Luo C, Poretsky R, Konstantinidis KT. 2011. Metagenomic insights into the evolution, function, and complexity of the planktonic microbial community of Lake Lanier, a temperate freshwater ecosystem. Appl Environ Microbiol 77:6000–6011.

10. Konstantinidis KT, DeLong EF. 2008. Genomic patterns of recombination, clonal divergence and environment in marine microbial populations. ISME J 2:1052–1065.

11. Olm MR, Crits-Christoph A, Diamond S, Lavy A, Matheus Carnevali PB, Banfield JF. 2020. Consistent metagenome-derived metrics verify and delineate bacterial species boundaries. mSystems 5:e00731–19.

12. Kim M, Oh HS, Park SC, Chun J. 2014. Towards a taxonomic coherence between average nucleotide identity and 16S rRNA gene sequence similarity for species demarcation of prokaryotes. Int J Syst Evol Microbiol 64:346–351.

13. Yarza P, Yilmaz P, Pruesse E, Glockner FO, Ludwig W, Schleifer KH, Whitman WB, Euzeby J, Amann R, Rossello-Mora R. 2014. Uniting the classification of cultured and uncultured bacteria and archaea using 16S rRNA gene sequences. Nat Rev Microbiol 12:635–645.

14. DeLong EF, Preston CM, Mincer T, Rich V, Hallam SJ, Frigaard N-U, Martinez A, Sullivan MB, Edwards R, Brito BR, Chisholm SW, Karl DM. 2006. Community genomics among stratified microbial assemblages in the ocean’s interior. Sience 311:486–503.

15. Sun DL, Jiang X, Wu QL, Zhou NY. 2013. Intragenomic heterogeneity of 16S rRNA genes causes overestimation of prokaryotic diversity. Appl Environ Microbiol 79:5962–5969.

16. Mende DR, Sunagawa S, Zeller G, Bork P. 2013. Accurate and universal delineation of prokaryotic species. Nat Methods 10:881–884.

17. Powell E, Ratnayeke N, Moran NA. 2016. Strain diversity and host specificity in a specialized gut symbiont of honeybees and bumblebees. Mol Ecol 25:4461–4471.

18. Raymann K, Bobay LM, Moran NA. 2018. Antibiotics reduce genetic diversity of core species in the honeybee gut microbiome. Mol Ecol 27:2057–2066.

19. Bobay LM, Wissel EF, Raymann K. 2020. Strain structure and dynamics revealed by targeted deep sequencing of the honey bee gut microbiome. mSphere 5:e00694–20.

20. Mukherjee S, Stamatis D, Bertsch J, Ovchinnikova G, Sundaramurthi JC, Lee J, Kandimalla M, Chen IA, Kyrpides NC, Reddy TBK. 2021. Genomes OnLine Database (GOLD) v.8: overview and updates. Nucleic Acids Res 49:D723–D733.

21. Baldaufnl SL, Baldaufnl AJ, Wenk-Siefert I, Doolittle WF. 2000. A kingdom-level phylogeny of Eukaryotes based on combined protein data. Sience 290:972–977.

22. Konstantinidis KT, Tiedje JM. 2005. Towards a genome-based taxonomy for prokaryotes. J Bacteriol 187:6258–6264.

23. Wu D, Jospin G, Eisen JA. 2013. Systematic identification of gene families for use as “markers” for phylogenetic and phylogeny-driven ecological studies of bacteria and archaea and their major subgroups. PLoS One 8:e77033.

24. Nayfach S, Rodriguez-Mueller B, Garud N, Pollard KS. 2016. An integrated metagenomics pipeline for strain profiling reveals novel patterns of bacterial transmission and biogeography. Genome Res 26:1612–1625.

25. Zheng H, Steele MI, Leonard SP, Motta EVS, Moran NA. 2018. Honey bees as models for gut microbiota research. Lab Anim (NY) 47:317–325.

26. Voulgari-Kokota A, McFrederick QS, Steffan-Dewenter I, Keller A. 2019. Drivers, diversity, and functions of the solitary-bee microbiota. Trends Microbiol 27:1034–1044.

27. Kwong WK, Medina LA, Koch H, Sing KW, Soh EJY, Ascher JS, Jaffé R, Moran NA. 2017. Dynamic microbiome evolution in social bees. Science Advances 3:e1600513.

28. Ellegaard KM, Suenami S, Miyazaki R, Engel P. 2020. Vast differences in strain-level diversity in the gut microbiota of two closely related honey bee species. Curr Biol 30:2520–2531 e2527.

29. Edgar RC. 2018. Updating the 97% identity threshold for 16S ribosomal RNA OTUs. Bioinformatics 34:2371–2375.

30. Martiny AC, Treseder K, Pusch G. 2013. Phylogenetic conservatism of functional traits in microorganisms. ISME J 7:830–838.

31. Langille MG, Zaneveld J, Caporaso JG, McDonald D, Knights D, Reyes JA, Clemente JC, Burkepile DE, Vega Thurber RL, Knight R, Beiko RG, Huttenhower C. 2013. Predictive functional profiling of microbial communities using 16S rRNA marker gene sequences. Nat Biotechnol 31:814–821.

32. Seemann T. 2014. Prokka: rapid prokaryotic genome annotation. Bioinformatics 30:2068–2069.

33. Page AJ, Cummins CA, Hunt M, Wong VK, Reuter S, Holden MT, Fookes M, Falush D, Keane JA, Parkhill J. 2015. Roary: rapid large-scale prokaryote pan genome analysis. Bioinformatics 31:3691–3693.

34. Katoh K, Standley DM. 2013. MAFFT multiple sequence alignment software version 7: improvements in performance and usability. Mol Biol Evol 30:772–780.

35. Stamatakis A. 2014. RAxML version 8: a tool for phylogenetic analysis and post-analysis of large phylogenies. Bioinformatics 30:1312–1313.

36. Yu G, Lam TT, Zhu H, Guan Y. 2018. Two Methods for mapping and visualizing associated data on phylogeny using ggtree. Mol Biol Evol 35:3041–3043.

37. Letunic I, Bork P. 2011. Interactive Tree Of Life v2: online annotation and display of phylogenetic trees made easy. Nucleic Acids Res 39:W475–W478.

38. Pritchard L, Glover RH, Humphris S, Elphinstone JG, Toth IK. 2016. Genomics and taxonomy in diagnostics for food security: soft-rotting enterobacterial plant pathogens. Analytical Methods 8:12–24.

39. Eren AM, Maignien L, Sul WJ, Murphy LG, Grim SL, Morrison HG, Sogin ML. 2013. Oligotyping: differentiating between closely related microbial taxa using 16S rRNA gene data. Methods Ecol Evol 4:1111–1119.

40. Madeira F, Park YM, Lee J, Buso N, Gur T, Madhusoodanan N, Basutkar P, Tivey ARN, Potter SC, Finn RD, Lopez R. 2019. The EMBL-EBI search and sequence analysis tools APIs in 2019. Nucleic Acids Res 47:W636–W641.

41. Kwong WK, Moran NA. 2013. Cultivation and characterization of the gut symbionts of honey bees and bumble bees: description of *Snodgrassella alvi* gen. nov., sp. nov., a member of the family *Neisseriaceae* of the *Betaproteobacteria*, and *Gilliamella apicola* gen. nov., sp. nov., a member of *Orbaceae* fam. nov., *Orbales* ord. nov., a sister taxon to the order *Enterobacteriales’* of the *Gammaproteobacteria*. Int J Syst Evol Microbiol 63:2008–2018.

42. Bolyen E, Rideout JR, Dillon MR, Bokulich NA, Abnet CC, Al-Ghalith GA, Alexander H, Alm EJ, Arumugam M, Asnicar F, Bai Y, Bisanz JE, Bittinger K, Brejnrod A, Brislawn CJ, Brown CT, Callahan BJ, Caraballo-Rodríguez AM, Chase J, Cope EK, Da Silva R, Diener C, Dorrestein PC, Douglas GM, Durall DM, Duvallet C, Edwardson CF, Ernst M, Estaki M, Fouquier J, Gauglitz JM, Gibbons SM, Gibson DL, Gonzalez A, Gorlick K, Guo J, Hillmann B, Holmes S, Holste H, Huttenhower C, Huttley GA, Janssen S, Jarmusch AK, Jiang L, Kaehler BD, Kang KB, Keefe CR, Keim P, Kelley ST, Knights D, Koester I, Kosciolek T, Kreps J, Langille MGI, Lee J, Ley R, Liu YX, Loftfield E, Lozupone C, Maher M, Marotz C, Martin BD, McDonald D, McIver LJ, Melnik AV, Metcalf JL, Morgan SC, Morton JT, Naimey AT, Navas-Molina JA, Nothias LF, Orchanian SB, Pearson T, Peoples SL, Petras D, Preuss ML, Pruesse E, Rasmussen LB, Rivers A, Robeson MS 2nd, Rosenthal P, Segata N, Shaffer M, Shiffer A, Sinha R, Song SJ, Spear JR, Swafford AD, Thompson LR, Torres PJ, Trinh P, Tripathi A, Turnbaugh PJ, Ul-Hasan S, van der Hooft JJJ, Vargas F, Vázquez-Baeza Y, Vogtmann E, von Hippel M, Walters W, Wan Y, Wang M, Warren J, Weber KC, Williamson CHD, Willis AD, Xu ZZ, Zaneveld JR, Zhang Y, Zhu Q, Knight R, Caporaso JG. 2019. Reproducible, interactive, scalable and extensible microbiome data science using QIIME 2. Nat Biotechnol 37:852–857.

43. Callahan BJ, McMurdie PJ, Rosen MJ, Han AW, Johnson AJ, Holmes SP. 2016. DADA2: High-resolution sample inference from Illumina amplicon data. Nat Methods 13:581–583.

44. Mirzajani A, Asharlous A, Kianpoor P, Jafarzadehpur E, Yekta A, Khabazkhoob M, Hashemi H. 2019. Repeatability of curvature measurements in central and paracentral corneal areas of keratoconus patients using Orbscan and Pentacam. J Curr Ophthalmol 31:382–386.

45. Soh EJY, Jaffé R, Ascher JS, Koch H, Medina LA, Moran NA, Kwong WK, Sing K-W. 2017. Dynamic microbiome evolution. Science Advances 3:e1600513.

46. Chen S, Zhou Y, Chen Y, Gu J. 2018. fastp: an ultra-fast all-in-one FASTQ preprocessor. Bioinformatics 34:i884–i890.

47. Rognes T, Flouri T, Nichols B, Quince C, Mahé F. 2016. VSEARCH: a versatile open source tool for metagenomics. PeerJ 4:e2584.

48. Zhang X, Li X, Su Q, Cao Q, Li C, Niu Q, Zheng H. 2019. A curated 16S rRNA reference database for the classification of honeybee and bumblebee gut microbiota. Biodiversity Science 27:557–566.

49. Li H, Durbin R. 2010. Fast and accurate long-read alignment with Burrows-Wheeler transform. Bioinformatics 26(5):589–595.

